# Regulation of lipid accumulation-induced ROS in myeloid-derived suppressor cells via targeting fatty-acid transport protein 2 enhanced anti-PD-L1 tumor immunotherapy

**DOI:** 10.1101/2020.09.28.316091

**Authors:** Adeleye Oluwatosin Adeshakin, Wan Liu, Funmilayo O. Adeshakin, Lukman O. Afolabi, Mengqi Zhang, Guizhong Zhang, Lulu Wang, Zhihuan Li, Lilong Lin, Dehong Yan, Xiaochun Wan

**Author notes:** Correspondence should be addressed to Dehong Yan or Xiaochun Wan.

## Abstract

Despite the remarkable success and efficacy of immune checkpoint blockade (ICB) therapy against the PD-1/PD-L1 axis, it induces sustained responses in a sizeable minority of cancer patients due to the activation of immunosuppressive factors such as myeloid-derived suppressor cells (MDSCs). Inhibiting the immunosuppressive function of MDSCs is critical for successful cancer ICB therapy. Interestingly, lipid metabolism is a crucial factor in modulating MDSCs function. Fatty acid transport protein 2 (FATP2) conferred the function of PMN-MDSCs in cancer via the upregulation of arachidonic acid metabolism. However, whether regulating lipid accumulation in MDSCs by targeting FATP2 could block MDSCs reactive oxygen species (ROS) production and enhance PD-L1 blockade-mediated tumor immunotherapy remains unexplored. Here we report that FATP2 regulated lipid accumulation, ROS, and immunosuppressive function of MDSCs in tumor-bearing mice. Tumor cells-derived granulocyte macrophage-colony stimulating factor (GM-CSF) induced FATP2 expression in MDSCs by activation of STAT3 signaling pathway. Pharmaceutical blockade of FATP2 expression in MDSCs by lipofermata decreased lipid accumulation, reduced ROS, blocked immunosuppressive activity, and consequently inhibited tumor growth. More importantly, lipofermata inhibition of FATP2 in MDSCs enhanced anti-PD-L1 tumor immunotherapy via the upregulation of CD107a and reduced PD-L1 expression on tumor-infiltrating CD8^+^ T-cells. Furthermore, the combination therapy blocked MDSC’s suppressive role on T-cells thereby enhanced T-cell’s ability for the production of IFN-γ. These findings indicate that FATP2 plays a key role in modulating lipid-induced ROS in MDSCs and targeting FATP2 in MDSCs provides a novel therapeutic approach to enhance anti-PD-L1 cancer immunotherapy.

**Research highlights:**

- FATP2 inhibition in MDSCs blocked ROS-mediated immunosuppressive function and promoted MDSCs differentiation to immune-stimulatory phenotype.
- GM-CSF mediated FATP2 expression via activation of STAT3 signaling leading to lipid accumulation-induced ROS in MDSCs.
- FATP2 inhibition enhanced anti-PD-L1 tumor immunotherapy by reducing MDSCs immunosuppressive activity in blocking T-cell’s ability to produce IFN-γ.
- Combination of anti-PD-L1 antibody and FATP2 inhibitor decreased MDSCs accumulation and PD-L1 surface expression in the spleens of LLC-bearing mice.
- Combined treatment of FATP2 inhibitor and PD-L1 blockade-mediated tumor immunotherapy enhanced tumor-infiltrating CD8^+^ T cells activation via increased CD107a and reduced PD-L1 surface expression.

## 1. Introduction

Immune checkpoint blockade (ICB) targeting programmed death receptor 1/ programmed death-ligand 1 (PD-1/PD-L1) pathway achieved clinical successes in the treatment of only a few tumors such as melanoma and non-small cell lung cancer [1]. However, metastatic melanoma patients who had initial responses to ICB developed resistance after about 2 years following the commencement of ICB treatment [2, 3]. The major cause of this resistance is the presence of immunosuppressive cells such as myeloid-derived suppressor cells (MDSCs), tumor-associated macrophages, tumor-associated dendritic cells, and regulatory T-cells in the tumor microenvironment. MDSCs contribute mainly to the formation of the pro-tumor immunosuppressive microenvironment and their levels could predict responses or resistance to ICB in cancer patients[4].

MDSCs are a heterogeneous population of immature myeloid cells involved in tumor progression, metastasis, and immune tolerance [5–8]. Phenotypically, these cells are similar to neutrophils and monocytes; however, biochemically and functionally, they are different from the aforementioned cell subsets [5]. In mice, MDSCs are characterized as cells co-expressing CD11b and Gr1 markers, which can be further categorized into two populations: polymorphonuclear (PMN)-MDSCs (CD11b^+^Ly6G^+^Ly6C^low^) and monocytic (M)-MDSCs (CD11b^+^Ly6G^−^Ly6C^high^) [8–10]. Functionally, MDSCs use different mechanisms such as induction of arginase 1 (Arg1), inducible nitric oxide synthase (iNOS), and reactive oxygen species (ROS) production to suppress anti-tumor immune responses [11]. Thus targeting the immunosuppressive function of MDSCs is critical for successful cancer ICB therapy.

Lipid metabolism is an important regulator of MDSCs function in tumors [9, 12–14] and could contribute to ROS-mediated immune suppression. Accumulation of lipids in cells could either be through fatty acid synthesis or exogenous uptake [15–17]. While other tumor-infiltrating myeloid cells such as dendritic cells and activated macrophages accumulate lipid through fatty acid synthesis [18, 19], MDSCs have been reported to prefer extrinsic fatty acids uptake with tumor-infiltrating MDSCs using fatty acid oxidation (FAO) as their energy sources [13, 20–22]. Our group was the first to show polyunsaturated fatty acid (PUFA) promoted MDSCs expansion and its immunosuppressive activity in tumors [12]. We reported that PUFA induced its effect through the activation of JAK/STAT3 signaling pathway while the inhibition of STAT3 activation in MDSCs almost completely abrogated its immunosuppressive function. Increased fatty acid uptake activated FAO in MDSCs while treatment with etomoxir, an FAO inhibitor, altered immune-inhibitory function, and enhanced chemotherapy [20]. Also, Al-Khami et al., highlighted the involvement of tumor-derived growth factors such as GM-CSF or G-CSF in inducing fatty acid uptake to reprogram MDSCs function in tumor milieu via either STAT3/STAT5 signaling pathways [13]. In another study, Vegilia et al., highlighted the role of GM-CSF in the upregulation of lipid metabolic gene via activation of STAT5 [21]. Collectively, these studies indicated lipid metabolism modulates the immunosuppressive function of MDSCs and possibly contributes to immunotherapy resistance in cancer patients.

Exogenous fatty acid uptake is mediated by active transporters which are members of the fatty acid transport proteins family (FATP; SLC27A) [23–25]. The FATP family comprises of six members referred to as FATP 1-6. It is found in the plasma membrane and intracellular organelles and possesses fatty acyl-CoA ligase activity [25, 26]. FATP is established to promote cellular fatty acid uptake in the liver, brain, heart, adipocytes, immune, and tumor cells [23, 27–29]. In a recent study, FATP2 was reported to be upregulated in PMN-MDSCs from mice and cancer patients, it was observed that FATP2 overexpression regulated the function of PMN-MDSCs in cancer via uptake of arachidonic acid to synthesize prostaglandin E2 (PGE2) [21]. The deletion of FATP2 in tumor-bearing mice reduced PMN-MDSCs activity and substantially inhibited tumor progression.

However, whether regulation of lipid accumulation in MDSCs by targeting FATP2 could control ROS activity and enhance PD-L1 blockade-mediated tumor immunotherapy remains unknown. Here we report a new role of FATP2 in regulating MDSCs function via lipid accumulation-induced ROS and targeting FATP2 in MDSCs promises a new approach in overcoming resistance to anti-PD-L1 therapy.

## 2. Materials and Methods

### 2.1. Reagent and Antibodies

DMEM, RPMI 1640, fetal bovine serum-albumin (FBS), penicillin-streptomycin (PS), Trypsin-EDTA, and Phosphate buffer saline (PBS) were purchased from Hyclone, USA. CFSE (Invitrogen), RIPA (Beyotime), 5X sample SDS buffer (GeneStar), Recombinant murine GM-CSF, and IL-6 (PeproTech). 5-bromo-5′-phenylspiro [3H-1, 3, 4-thiadiazole-2, 3′-indoline] −2-one (Lipofermata) (Cayman), InVivoMab anti-mouse PD-L1 antibody (clone 10F.9G2, BioXCell), Stattic (MedChem Express), BD Cytofix/Cytoperm kit (BD Bioscience). The free fatty acids (linoleic, Oleic, α-linolenic, and stearic acid), 4, 6-diamidino-2-phenylindole (DAPI), Brefeldin A. The following molecular probes: BODIPY (493/503), BODIPY FLC16, Chloromethyl-2,7-Dichlorofl-fluorescein Diacetate (CM-H_2_DCFDA), mitotracker green, mitotracker red, and mitosox red as well as Lipofectamine 3000 were obtained from Life Technologies, Invitrogen. The antibodies against phospho-STAT3, actin, and HRP conjugated secondary antibodies against mouse and rabbit were from Cell Signaling Technology Inc.; FATP4 and STAT3 were Santa Cruz while FATP2 was from Thermofischer scientific. The following fluorescein-conjugated anti-mouse antibodies: CD11b-PE, Gr1-APC, Ly6C-FITC, Ly6G-APC, CD11c-PerCP-Cy5.5, F4/80-APC, CD3-PE, CD4-PerCP-Cy5.5, CD8a-PE, CD19-FITC, NK1.1-APC, CD107a - PerCP-Cy5.5, CD69-PE, PD-L1-PE (clone MIH5), GM-CSF-PE (clone MP1-22E9), IL-6-PE, IL-10-PE, IFN-γ-PE, TNF-α-PE; purified CD3 and CD28; CD45 nanobeads, CD11b biotin, Gr1 biotin, anti-biotin beads, purified anti-mouse GM-CSF antibody (clone MP1-31G6) were from Biolegend, CA, USA.

### 2.2. Cell culturing

B16F10 and LLC cells were obtained from cell bank, Chinese Academy of Sciences. It was sub-cultured using DMEM with 10% FBS and 1% antibiotics, grown in a 37 °C, 5% incubator. The medium was changed every 2-3 days and cell passage was done once the cell grew to about 90% confluence.

### 2.3. Preparation of tumor explanted supernatant (TES)

B16F10 or LLC tumor tissue from C57BL/6 mice without any ulceration, grown to about 2 cm diameter was carefully removed under a sterile condition after sacrificing the mice. Cells were bathed in 70% ethanol for 30 seconds and digested with Collagenase I, II, IV, and hyaluronidase for 1 hour at 37°C. The digested tissue was passed through a 70 µm mesh to obtain a single cell suspension and the supernatant was centrifuged at 1260 g for 10 minutes. The cell pellet was washed with PBS and re-suspended in RPMI containing 3% FBS and 1% PS. Cells were cultured at 1×10^7^ cells/ml under 37°C, 5% CO_2_ for 18-24 hours and cell-free supernatant collected using a 0.22 µm filter membrane for immediate use or storage at –80°C.

### 2.4 Generation of MDSC in-vitro

Bone marrow cells were isolated from wild-type (WT) naive and tumor-bearing C57BL/6 mice. Ammonium-chloride-potassium (ACK) buffer was used to lyse the red blood cells of isolated cells. GM-CSF (40 ng/ml) were used to generate MDSCs from cultured bone marrow cells in RPMI with 10% FBS and 1% antibiotics for 3 days and incubated at 37°C, 5% CO_2_. The generated MDSCs were subjected to further experimental procedures.

### 2.5 Cell viability assay

50,000 CD11b^+^Gr1^+^ MDSCs from the bone marrow of naive mice stimulated with GM-CSF plus IL-6 or 10,000 cells of B16F10 and LLC cell lines were plated in 96 well plates for 24 hours before treatment. On the following day, cells were treated with varying concentrations of lipofermata for 3 days while PBS served as the vehicle. Cell viability was assayed in triplicates using CellTiter 96 cell proliferation assay kit (Promega Madison, USA) according to the manufacturer’s instruction. Relatively viable cells were determined by subtracting the optical density at 630 nm (background) from 490 nm.

### 2.6 Flow cytometry

A single-cell suspension of the spleen, bone marrow, or tumor in PBS containing 2% FBS were labeled with the appropriate fluorescein-conjugated anti-mouse antibodies or molecular probes. The percentages of various immune cells, lipid content, and oxidative stress markers were evaluated by BD FACS Canto^TM^ II flow cytometer (BD Bioscience). The acquired data were processed using flowJo software (version 7.6). Purified MDSCs from spleen and bone marrow cells were isolated using a mojosort magnet followed by cell surface staining with CD11b and Gr1 monoclonal antibodies for cell sorting on BD FACS Aria III cell sorter (BD Biosciences). Similarly, CD8^+^ T cells were obtained from the spleen of naive mice following antibody labeling and sorted on flow cytometer BD FACS Aria III cell sorter (BD Biosciences).

### 2.7 Lipid accumulation determination

To determine lipid accumulation, cells previously incubated with specific antibodies against myeloid cells were washed twice with PBS and re-suspended in 500µl BODIPY 493/503 at 500 µg/ml for 15 minutes in the dark at room temperature. Cells were then washed twice with PBS and re-suspended in 300µl of PBS for flow cytometry analyses.

### 2.8 Fatty-acid uptake analyses

Cells were stained with surface markers followed with fluorescently labeled BODIPY FLC16 at 10µg/ml which incorporates palmitic acid into the cells. Samples were incubated for 30 minutes protected from light at 37°C before analyses on the flow cytometer.

### 2.9 Measurement of total reactive oxygen species (ROS), mitochondrial ROS, superoxide anion production, and mitochondrial mass

Cells suspension previously incubated with CD11b and Gr1 antibodies were labeled with the following molecular probes: mitotracker green to determine mitochondrial mass, mitosox red to evaluate mitochondrial superoxide anion production, mitotracker red to measure mitochondrial ROS, and chloromethyl-2, 7-dichlorofluorescein diacetate (CM-H_2_DCFDA) to determine total ROS production. Cells were incubated for 30 mins protected from light at 37°C before analyses on the flow cytometer.

### 2.10. RT-qPCR gene expression analyses

*In-vivo* and *in-vitro* generated CD11b^+^ Gr1^+^ MDSCs were sorted on flow cytometry machine and cells pelleted for RNA isolation using RNAiso plus (Takara bio. Inc., Japan). The extracted RNA was converted to cDNA with Takara kit (Takara bio. Inc., Japan). The transcript level of different genes of interest was evaluated via qPCR machine using SYBR green master mix (MedChem Express, USA) and detected on CFX96^TM^ Real-Time System C1000 Touch Thermal cycler (BIO-RAD); Conditions: (1) 95°C, 5 minutes; (2) 95°C, 15 seconds; (3) 60°C, 45 seconds; (4) 65°C, 5 seconds; (5) 95°C, 50 seconds; 40 cycles. Amplification of endogenous actin served as the housekeeping gene. The primers sequence are listed in Supplementary Table S1.

### 2.11. Lentivirus-mediated gene silencing

HEK293T cells (ATTC) were transfected with mouse STAT3 shRNA and scrambled control shRNA plasmids (sequence is shown in Table S2) using lipofectamine 3000 according to the manufacturer’s instructions. The virus supernatant was collected 48 hours post-transfection, filtered, concentrated by ultracentrifugation, and titrated by limiting dilution assay. Bone-marrow derived MDSCs were infected with STAT3 shRNA lentivirus for 48 hours and cells harvested to determine knockdown efficiency via measuring the mRNA expression level of STAT3 using RT-qPCR.

### 2.12. Immunoblotting

Purified MDSCs from spleen or bone marrow that received various treatments were lysed on ice using RIPA lysis buffer. Cell lysates were centrifuged at 12,000g, 4°C for 10 minutes, and the supernatant was collected to determine the protein concentration using the Pierce^TM^ BCA protein Assay kit (Thermo Fisher Scientific). 1x sample SDS buffer was added to the supernatant for electrophoresis. 30-50µg of protein per lane alongside a prestained molecular weight protein marker was separated on an 8% SDS PAGE gel prepared from SDS-PAGE kit (Beyotime, China) and electrotransferred to immunoblot PVDF membrane (BIO-RAD, CA, USA) for protein blotting. After blocking of the membrane in western quick block kit (Beyotime, China) or 5% non-fat dry milk for at least 1 hour, it was incubated in primary antibodies against STAT3, p-STAT3, FATP2, and FATP4 with gentle agitation overnight at 4°C. Actin was used as the housekeeping protein. HRP-conjugated secondary antibodies were used to incubate the membrane for an hour followed by protein detection with enhanced chemiluminescence (ECL) western blotting substrate and viewed on Amersham Imager 600 (GE Healthcare). Uncropped immunoblot images are shown in Figure S8.

### 2.13. Cytokine Detection

Cell supernatants were obtained from bone-marrow-derived CD11b^+^Gr1^+^ MDSCs previously incubated with fatty acids, TES or TES-lipofermata, and co-cultured with CD3^+^ T-cells from the splenocytes of naive mice for 36 hours. CD3^+^ T-cells was stimulated with purified anti-CD3 (1 µg/ml) and anti-CD28 (2 µg/ml) antibodies. IFN-γ and TNFα levels were determined using the BD cytometric bead array (CBA) cell signaling master buffer kit (San Jose, CA, US). ELISA was used to determine the release of GM-CSF or IL-6 from MDSCs supernatant and TES. Similarly, purified MDSCs from bone-marrow or spleen of LLC or B16F10 mice treated with lipofermata, anti-PD-L1 antibody, or combination of lipofermata and anti-PD-L1 as well as control mice were co-cultured with CD3^+^ T-cell from the splenocytes of naive mice for 36 hours. Brefeldin A (10µg/ml) was added to the cells and mixed thoroughly, it was later kept at 37°C, 5% CO_2_ for 3 hours before cell surface staining. Cell supernatant was used for CBA analysis while cell pellet was used for intracellular flow cytometry analysis of IFN-γ or TNFα. Cell pellets were then stained for surface marker (CD3), fixed with BD Cytofix/Cytoperm kit, and stained for intracellular cytokines (IFN-γ, TNFα).

### 2.14. Liquid chromatography-electrospray ionization-mass spectrophotometry (LC-ESI-MS)

CD11b^+^ Gr1^+^ flow cytometry sorted MDSCs from the spleen of naive and B16F10 tumor-bearing; bone marrow-derived MDSCs from naive mice induced with or without TES; spleen of B16F10 tumor-bearing mice with or without FATP2 inhibition were resuspended in 200ul of PBS and lipid was extracted according to a previously described method [30]. 1.5ml of methanol was added to the samples and vortexed vigorously for 30 seconds. 5ml of MTBE was added to the vortexed mixture and incubated for 1 hour on a shaker at room temperature. Phase separation was carried out by adding 1.25ml of MS-grade water to the above mixture. The samples were incubated for 10 minutes at room temperature and then centrifuged at 1000g for another 10 minutes. The upper phase was collected and placed in a new tube while the lower phase was re-extracted with 2 ml of the solvent mixture - MTBE/methanol/water (10:3:2.5, v/v/v). The upper phase was collected and added to the previously collected upper phase. The combined organic phase was dried in a vacuum centrifuge overnight. The vacuum dried samples were reconstituted in ACN/IPA/H_2_O (65:30:5, v/v/v) containing 5mM ammonium acetate, and 5μL was injected into the LC-ESI-MS system to identify and quantify different lipid species present in MDSCs.

A reversed-phase BEH C8 column (2.1mm 100mm, 1.7μm, Waters, Milford, MA, U.S.A.) was used for the chromatographic separation of lipids. Mobile phases A and B were ACN/H_2_O (60:40, v/v) and IPA/ACN (90:10, v/v) respectively, both containing 10mM ammonium acetate. The flow rate was 0.26 mL/minutes while the column temperature was at 55 °C. The elution started with 68% mobile phase A (ACN: H_2_O = 6:4, 10mM ammonium acetate) and 32% mobile phase B (IPA: ACN = 9:1, 10mM ammonium acetate) and maintained for 1.5 minutes. Mobile phase B was then linearly increased to 85% for 10 minutes and further to 97% in the next 0.1 minutes followed by maintenance for 1.5 minutes. Afterward, it was decreased to 32% B in 0.1 minutes and kept for 2 minutes till the next injection. The temperature of the sample manager was set at 10 °C.

The mass spectrometer was operated with a capillary voltage of 3.5 kV in positive mode and 3.0 kV in negative mode. The capillary temperature was set at 300°C. Sheath gas flow rate and aux gas flow rate were set at 30 and 10 (in arbitrary units) respectively. Aux gas heater temperature was 310 °C while the S-lens RF level was 50.0. The resolutions of 70000 and 17500 were set for full scan MS and data-dependent MS/MS (ddMS2) in both modes. AGC target and maximum IT were 3×10^6^ ions capacity and 200ms in full-scan MS settings while their values were 1×10^5^ ions capacity and 50ms in ddMS2 settings. The Top N (N, the number of topmost abundant ions for fragmentation) was set to 10. The normalized collision energy (NCE) was set 20, 35, 70eV, respectively while the scan range was set at m/z 133.4−2000.

### 2.15. In-vivo experiments

All animal experiments were approved by the ‘Animal Care and Use Committee’ of the Shenzhen Institutes of Advanced Technology under the protocol SIAT-IACUC-190403-YYS-YXL-A0732. The mice were maintained in the animal facilities of Shenzhen Institutes of Advanced Technology, Chinese Academy of Sciences, under pathogen-free conditions. C57BL/6 and Balb/C nude mice (6–8-wk-old females) were obtained from Guangdong Province Animal Care Facilities. To establish subcutaneous tumors in C57BL/6 or Balb/c nude mice, 1×10^6^ B16F10 or LLC cells per mouse on the lower right flank were injected subcutaneously into the mice. Tumor size and body weights of the mice were recorded every other day starting from the point when tumor growth was palpable to about 3 weeks or more before mice were sacrificed. Tumor volume was measured using a digital caliper and calculated by using the formula [(length) x (width) ^2^ /2]. Mice were treated intraperitoneally with 2.5mg/kg lipofermata daily and 200µg of anti-PD-L1 antibody every three days from the first day of treatment while PBS was used as a vehicle in the control group. Spleen, bone marrow, and tumor were harvested from the mice at the end of the study. A single-cell suspension was prepared from the spleen, bone marrow cells, or tumor for further analysis. Also, weight and images of harvested tumor and spleen were taken.

### 2.16. Statistical analyses

Statistical analyses were performed using Prism 8.4.2 software (Graphpad Software, CA, USA). All data are presented as mean ± standard error of the mean (SEM), and P < 0.05 was considered significant. Each experiment was conducted at least three times unless otherwise indicated. Data analysis was performed by either student t-test, one-way, or two-way ANOVA with Tukey’s post-test. In figures, asterisks were used as follows: *, P ≤ 0.05; **, P ≤ 0.01; ***, P ≤ 0.001; and ****, P ≤ 0.0001.

## 3. Results

### 3.1 FATP2 regulates ROS-mediated immunosuppressive function of MDSCs

In the tumor microenvironment, MDSCs accumulate lipids via exogenous fatty acid uptake leading to enhanced mitochondrial function and activation of ROS, thus promoting the immunosuppressive function of MDSCs [11, 12, 20]. In our *in-vitro* experiments, we observed that TES enhanced mitochondrial function and ROS in MDSCs as shown by increased generation of total ROS (Figure 1A and S1A), mitochondrial ROS (Figure 1B and S1A), superoxide anion (Figure 1C and S1A) and mitochondrial mass (Figure 1D and S1A) compared to the control medium. Similar to the *in-vitro* observation, we noticed spleen MDSCs from B16F10-bearing mice increased total ROS, mitochondrial ROS, and mitochondrial mass compared to spleen CD11b^+^Gr1^+^ cells from tumor-free mice (Figure 1E-G and S1B). Interestingly, tumor MDSCs generated higher ROS compared to spleen and bone marrow MDSCs from B16F10-bearing mice (Figure 1E and S1B). More importantly, unsaturated fatty acids (linoleic acid-LA, oleic acid-OA, and α-linolenic acid-ALA) increased ROS generation of *in-vitro* derived MDSCs compared to the control group and saturated fatty acid (stearic acid-SA) (Figure 1H). Furthermore, cytometric bead array assay showed that unsaturated fatty acids (LA, OA, ALA), not SA induced MDSCs suppressed T cell’s ability to produce IFN-γ and TNFα compared to the control (Figure 1I). Collectively, these data show that tumor-derived unsaturated fatty acids enhanced mitochondrial function and activated ROS, thus induced the immunosuppressive activity of MDSCs.

**Figure 1:**
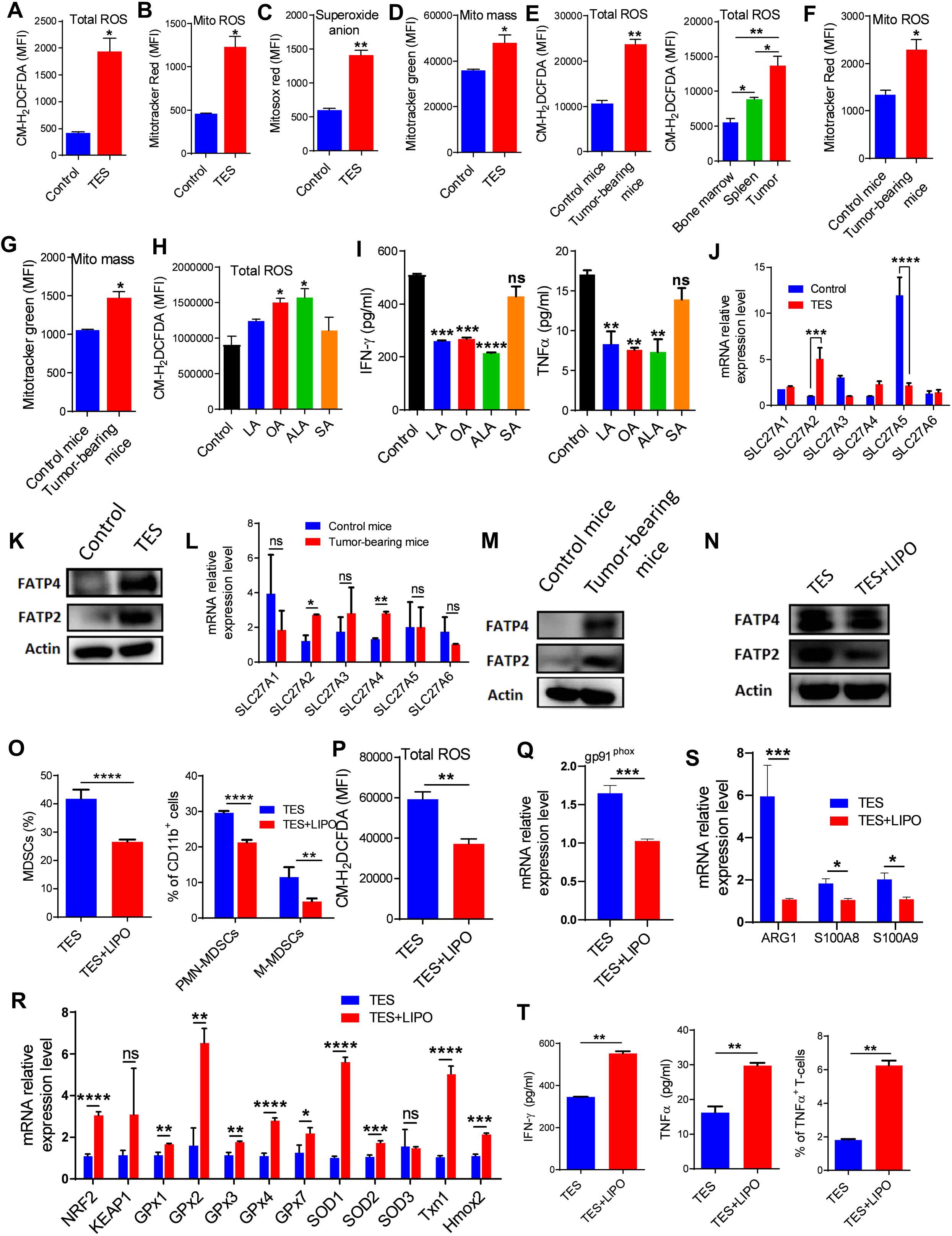
FATP2 regulates ROS-mediated immunosuppressive function of MDSCs. (A-D) Total ROS (A), mitochondrial ROS (B), superoxide anion (C), and mitochondrial mass (D) of CD11b^+^Gr1^+^ MDSCs derived from bone marrow cells cultured in the presence of control medium or TES. (E-G) Total ROS (E, left), mitochondrial ROS (F), mitochondrial mass (G) from spleen CD11b^+^Gr1^+^ cells from control tumor-free mice and spleen CD11b^+^Gr1^+^ MDSCs of B16F10 tumor-bearing mice. Total ROS of CD11b^+^Gr1^+^ MDSCs from the spleens, bone marrows, or tumor of B16F10 tumor-bearing mice (E, right). (H) Total ROS of CD11b^+^Gr1^+^ MDSCs generated from bone-marrow cells treated with medium, LA, OA, ALA, and SA for 48 hours. (I) The concentrations of IFN-γ and TNFα released by T cells as determined by CBA (left and right) in the co-culture of MDSCs and CD3^+^ T cells at ratio 1:2, respectively. CD3^+^ T cells were activated with the anti-CD3/CD28 purified antibody for 36 hours before testing. (J) RT-qPCR analyses of fatty acid transport proteins (SLC27A1-6/FATP1-6) in MDSCs generated from bone marrow cells cultured in the presence or absence of TES for 48 hours. (K) Immunoblotting to determine protein levels of FATP2 and FATP4 in MDSCs generated from bone marrow cells cultured in the presence or absence of TES for 48 hours. (L) RT-qPCR analyses of fatty acid transport proteins (SLC27A1-6/FATP1-6) in MDSCs from spleen CD11b^+^Gr1^+^ cells of tumor-free mice and spleen CD11b^+^Gr1^+^ MDSCs of B16F10-bearing mice. (M) Immunoblot analysis of FATP2 and FATP4 protein levels in MDSCs from spleen CD11b^+^Gr1^+^ cells of tumor-free mice and spleen CD11b^+^Gr1^+^ MDSCs of B16F10-bearing mice. (N) Expression levels of FATP2 and FATP4 as determined by immunoblotting in MDSCs from GM-CSF-induced bone marrow cells treated with TES in the presence or absence of lipofermata for 48 hours. (O) Proportion of total MDSCs (CD11b^+^Gr1^+^) (left), PMN-MDSCs (CD11b^+^Ly6G^+^Ly6C^low^) and M-MDSCs (CD11b^+^Ly6G^−^Ly6C^high^) (right) in GM-CSF-induced bone marrow cells-treated TES with or without lipofermata for 48 hours. (P) Total ROS in MDSCs from GM-CSF-induced bone marrow cells treated TES with or without lipofermata for 48 hours. (Q-S) RT-qPCR analysis for the expression of gp91^phox^ (Q), anti-oxidative genes (R), and immunosuppressive signatures (S) in MDSCs from GM-CSF-induced bone marrow cells treated TES with and without lipofermata for 48 hours. (T) The concentrations of IFN-γ and TNFα released by T cells was determined via CBA (left and middle) and percentage of TNFα^+^ T cells by intracellular flow cytometry staining (right) in the co-culture of MDSCs and CD3^+^ T cells at ratio 1:2, respectively. MDSCs were incubated in the presence of B16F10-TES and treated with or without lipofermata. CD3^+^ T cells were activated with the anti-CD3/CD28 purified antibody for 36 hours before testing. Data are shown as representative of mean ± SEM in triplicate samples from three independent experiments. Unpaired student’s t-test or one-way ANOVA: *p < 0.05, **p < 0.01, ***p < 0.001, ****p < 0.0001, ns, no significant difference.

Exogenous fatty acid uptake is mediated by fatty acid transport proteins (FATP) family [23–25]. Next, we asked which FATPs may be responsible for the increased ROS by MDSCs in tumor. Our RT-qPCR analysis revealed that tumor upregulated SLC27A2 (FATP2) and SLC27A4 (FATP4), downregulated SLC27A3 (FATP3) and SLC27A5 (FATP5), and did not affect SLC27A1 (FATP1) and SLC27A6 (FATP6) expression of *in-vitro* generated CD11b^+^Gr1^+^ MDSCs compared to the control medium (Figure 1J). By western blot analysis, we also detected that the protein expressions of FATP2 and FATP4 in TES-treated MDSCs were increased compared with control MDSCs (Figure 1K). Similarly, Spleen MDSCs of tumor-bearing mice showed upregulation in the mRNA expressions for FATP2 and FATP4 (Figure 1L) and protein expressions of FATP2 and FATP4 (Figure 1M) compared to spleen CD11b^+^Gr1^+^ cells of control tumor-free mice. However, no difference in the SLC27A1, SLC27A3, SLC27A5, and SLC27A6 mRNA expressions between spleen MDSCs of tumor-bearing mice and spleen CD11b^+^Gr1^+^ cells from control tumor-free mice (Figure 1L).

Considering the increased expression of FATP2 and FATP4 in MDSCs, we therapeutically targeted FATPs with a small molecule inhibitor, lipofermata. To establish if lipofermata targeted both FATP2 and FATP4 in MDSCs, we treated TES-induced MDSCs with lipofermata and then checked their protein expression by western blot analysis. Our results showed that lipofermata specifically inhibited FATP2 and not FATP4 expression in TES-induced MDSCs (Figure 1N). Also, lipofermata blocked the accumulation of B16F10-TES-induced MDSCs (Figure 1O and S1C). Importantly, our data showed that lipofermata decreased both PMN-MDSCs and M-MDSCs accumulation *in-vitro* (Figure 1O and S1C). To further investigate if FATP2 regulated ROS in MDSCs, we checked ROS levels and its regulatory genes expression via inhibition of FATP2 by lipofermata in TES-induced MDSCs. Interestingly, blockade of FATP2 by lipofermata reduced ROS generation of *in-vitro* TES-derived MDSCs (Figure 1P). More so, we observed a reduction in mRNA expression of *gp91^phox^*, a subunit of NADPH oxidase (NOX2) mediating ROS production in MDSCs (Figure 1Q). Besides, lipofermata upregulated the mRNA expression of anti-oxidative genes such as NRF2, GPx1-4, GPx7, SOD1-2, Txn1, and Hmox2, but did not change KEAP and SOD3 expression in TES-induced MDSCs compared to the untreated group (Figure 1R). Also, lipofermata downregulated the expression of immunosuppressive signature genes such as ARG1, S100A8, and S100A9 in TES-induced MDSCs (Figure 1S). Furthermore, inhibiting FATP2 decreased the suppressive role of MDSCs leading to enhanced CD3^+^ T cells to release more IFN-γ (left), TNFα (middle), and increase the percentages of TNFα^+^ CD3^+^ T cells (right) (Figure 1T and S1D). Our findings indicate that FATP2 regulates ROS-mediated suppressive function of MDSCs.

### 3.2 GM-CSF triggers FATP2 expression leading to lipid accumulation-induced ROS in MDSCs

Previous study showed tumor-derived cytokines could contribute to lipid accumulation in MDSCs [13]. We next investigated which could be the major tumor-derived cytokine contributing to lipid accumulation-induced ROS of MDSCs in the tumor environment. Using RT-qPCR, we evaluated the relative expression of GM-CSF, G-CSF, M-CSF, IL-6, TNFα, TGF-β, and IL-10 from CD11b^+^Gr1^+^ MDSCs in the bone marrows of tumor-bearing mice and B16F10 cell line. Our data revealed an upregulation of GM-CSF, G-CSF and M-CSF, and downregulation of IL-6, TNFα, and TGF-β in B16F10 cells compared to MDSCs (Figure 2A). However, we observed no significant difference in IL-10 expression (Figure 2A). It is well established that GM-CSF promotes the differentiation and accumulation of MDSCs in-vitro [31, 32] while IL-6 and IL-10 are critical to its suppressive function [33]. Next, we evaluated the percentages of GM-CSF, IL-6, and IL-10 positive cells using flow cytometry. Our observation shows that percentages of GM-CSF^+^ not IL-6^+^ tumor cells were higher compared to the percentages of spleen MDSCs (Figure 2B). The percentages of IL-10^+^ tumor cells were slightly higher than the percentages of spleen MDSCs (Figure 2B). Also, spleen MDSCs supernatant and B16F10 or LLC derived-TES confirmed the elevated release of GM-CSF in tumor cells as determined by ELISA (Figure 2C). Furthermore, we asked if LA, OA, ALA, and SA directly induce GM-CSF or IL-6 expression in MDSCs. Unexpectedly, the data reveals that fatty acids did not affect the percentage of GM-CSF^+^ MDSCs while LA, OA, and ALA increased expression of IL-6 in bone-marrow-derived MDSCs as detected by flow cytometry and ELISA (Figures S2A-C). We then investigated if GM-CSF or IL-6 modulates the lipid accumulation of MDSCs. Freshly harvested bone marrow cells from C57BL/6 mice were cultured in the presence of GM-CSF, IL-6, or both cytokines at varying concentrations for 3 days. Cells were harvested to evaluate the lipid level. Flow cytometry analysis showed that stimulation with either GM-CSF or IL-6 increased lipid accumulation of MDSCs in a dose-dependent manner compared to the unstimulated cells (Figure 2D). On the contrary, it was observed that only GM-CSF, not IL-6, enhanced the ability of MDSCs for the uptake of fluorescently labeled palmitic acid (BODIPY FLC16) compared to the unstimulated cell (Figure 2E). Next, we investigated if GM-CSF and IL-6 had a synergistic effect on lipid accumulation or uptake. Surprisingly, the combination of GM-CSF and IL-6 had no synergistic effect on lipid accumulation or uptake of MDSCs compared to GM-CSF only treated MDSCs (Figures 2F-G). These results suggest that GM-CSF is the key cytokine involved in lipid accumulation of MDSCs.

**Figure 2:**
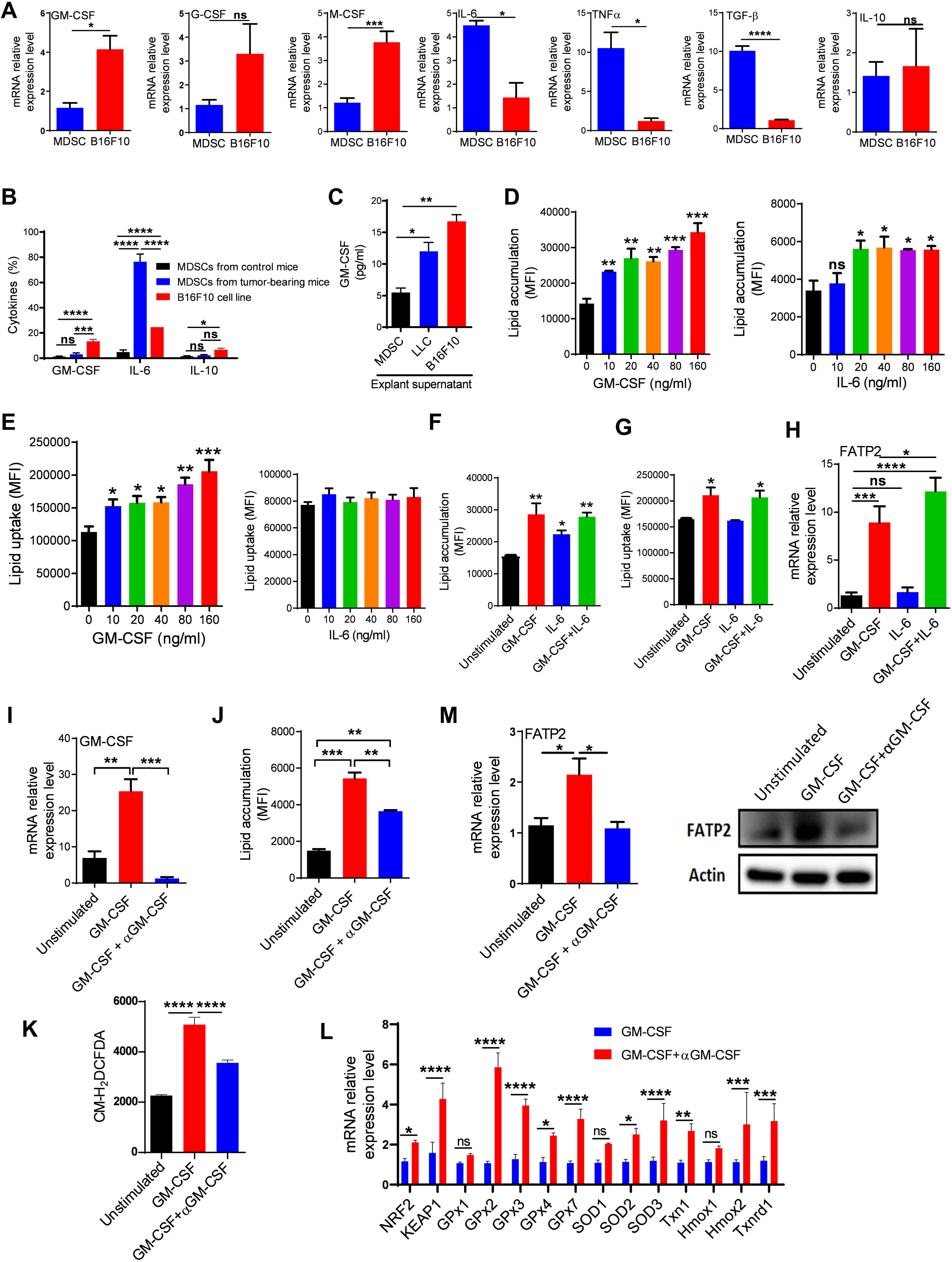
GM-CSF triggers FATP2 expression leading to lipid accumulation-induced ROS in MDSCs. (A) RT-qPCR analyses of GM-CSF, G-CSF, M-CSF, IL-6, TNFα, TGF-β, and IL-10 mRNA relative expression in bone marrow-derived MDSCs from B16F10 tumor-bearing mice and B16F10 cell line. (B) Percentage of GM-CSF^+^, IL-6^+^, and IL-10^+^ in spleen CD11b^+^Gr1^+^ cells of tumor-free or spleen CD11b^+^Gr1^+^ MDSCs from B16F10 tumor-bearing mice or B16F10 cell line as determined by flow cytometry. (C) GM-CSF concentration in spleen MDSCs supernatant or TES from B16F10 or LLC tumor-bearing mice. (D) Lipid accumulation in MDSCs from bone-marrow cells stimulated with varying concentrations of GM-CSF (left) and IL-6 (right) compared to unstimulated MDSCs for 72 hours. (E) Exogenous lipid uptake levels in MDSCs from bone-marrow cells stimulated with indicated concentrations of GM-CSF (left) and IL-6 (right) for 72 hours compared to unstimulated MDSCs. (F-H) MDSCs from bone marrow cells *in vitro* stimulated with GM-CSF, IL-6, or both GM-CSF and IL-6 for 72 hours compared to unstimulated MDSCs. Mean fluorescence intensity (MFI) for lipid accumulation by flow cytometry analysis (F), MFI for exogenous lipid uptake by flow cytometry analysis (G), and RT-qPCR analyses for FATP2 mRNA expression of MDSCs (H). (I-M) mRNA expression for MDSCs from bone marrow cells *in vitro* stimulated with GM-CSF only or GM-CSF and anti-GM-CSF antibody (5µg/ml) compared to unstimulated MDSCs for 24 hours, as determined by RT-qPCR analysis (I), Flow cytometry analysis for lipid accumulation (J), Total ROS level as determined by flow cytometry (K), mRNA level of anti-oxidative genes as determined by RT-qPCR (L), FATP2 mRNA expression determined by RT-qPCR analysis (M, left) and protein expression determined by immunoblot analysis (M, right) of MDSCs. MDSCs were CD11b^+^Gr1^+^ cells; Data are shown as representative of mean ± SEM in triplicate samples from three independent experiments. Unpaired student’s t-test or one-way ANOVA: *p < 0.05, **p < 0.01, ***p < 0.001, ****p < 0.0001, ns, no significant difference.

Next, we asked if GM-CSF could induce ROS and triggers FATP2 expression leading to lipid accumulation in MDSCs. Interestingly, GM-CSF upregulated the expression of FATP2 while IL-6 did not affect it compared to the unstimulated cell; although the combination of GM-CSF and IL-6 synergistically enhanced FATP2 expression (Figure 2H). On the other hand, we observed significant downregulation of FATP4 expression in MDSCs stimulated with either GM-CSF or IL-6 as well as their combination (Figure S2D). We moved on to block the GM-CSF signaling in MDSCs by anti-GM-CSF antibody and we confirmed with RT-qPCR that anti-GM-CSF antibody blocked GM-CSF signaling in MDSCs (Figure 2I). Interestingly, blocking GM-CSF signaling decreased lipid accumulation (Figure 2J), reduced ROS production (Figure 2K), upregulated antioxidant genes (Figure 2L), downregulated FATP2 gene and protein expression (Figure 2M) in MDSCs compared with GM-CSF-induced MDSCs. However, the blockade of GM-CSF signaling did not affect FATP4 expression (Figure S2E). Therefore, we propose that GM-CSF is the major cytokine involved in the upregulation of FATP2 expression leading to lipid accumulation-induced ROS in MDSCs.

### 3.3 GM-CSF induces FATP2 expression by STAT3 activation in tumor MDSCs

Our previous work demonstrated the involvement of STAT3 signaling in mediating the suppressive effect of polyunsaturated fatty acid-induced MDSCs [12]. Also, GM-CSF has been reported to promote the activation of STAT3 signaling in MDSCs [34, 35]. To explore if GM-CSF controls FATP2 expression through activation of STAT3 signaling, we used a small molecule inhibitor stattic, to block STAT3 activation in *in-vitro* GM-CSF-induced bone marrow MDSCs. Firstly, we confirmed via western blot that GM-CSF activated STAT3 signaling by upregulation of phospho-STAT3 and stattic did inhibit GM-CSF-induced STAT3 activation in MDSCs (Figure 3A). Next, we observed that blockade of STAT3 signaling by stattic resulted in a significant reduction in ROS production (Figure 3B), and significantly inhibited GM-CSF-induced FATP2 expression in MDSCs (Figure 3C). According to Figure 2C, GM-CSF was mainly derived from TES, so we wanted to see if inhibition of STAT3 signaling in TES-induced MDSCs by shRNA-mediated knockdown or stattic could regulate FATP2 expression. RT-qPCR analysis showed that STAT3 shRNA downregulated STAT3 and FATP2 mRNA expression in TES-treated MDSCs compared to control shRNA-treated TES-induced MDSCs (Figures 3D-E). Interestingly, blockade of STAT3 activation by stattic dramatically downregulate FATP2 mRNA and protein expression (Figures 3F-G), and also reduced lipid accumulation, lipid uptake, and ROS production in TES-induced MDSCs compared to untreated TES-induced MDSCs (Figures 3H-J). But we noticed that inhibiting STAT3 activation in control MDSCs slightly reduced lipid accumulation, lipid uptake, and ROS production (Figures 3H-J). Collectively, these suggest that GM-CSF induces FATP2 expression by STAT3 activation in tumor MDSCs.

**Figure 3:**
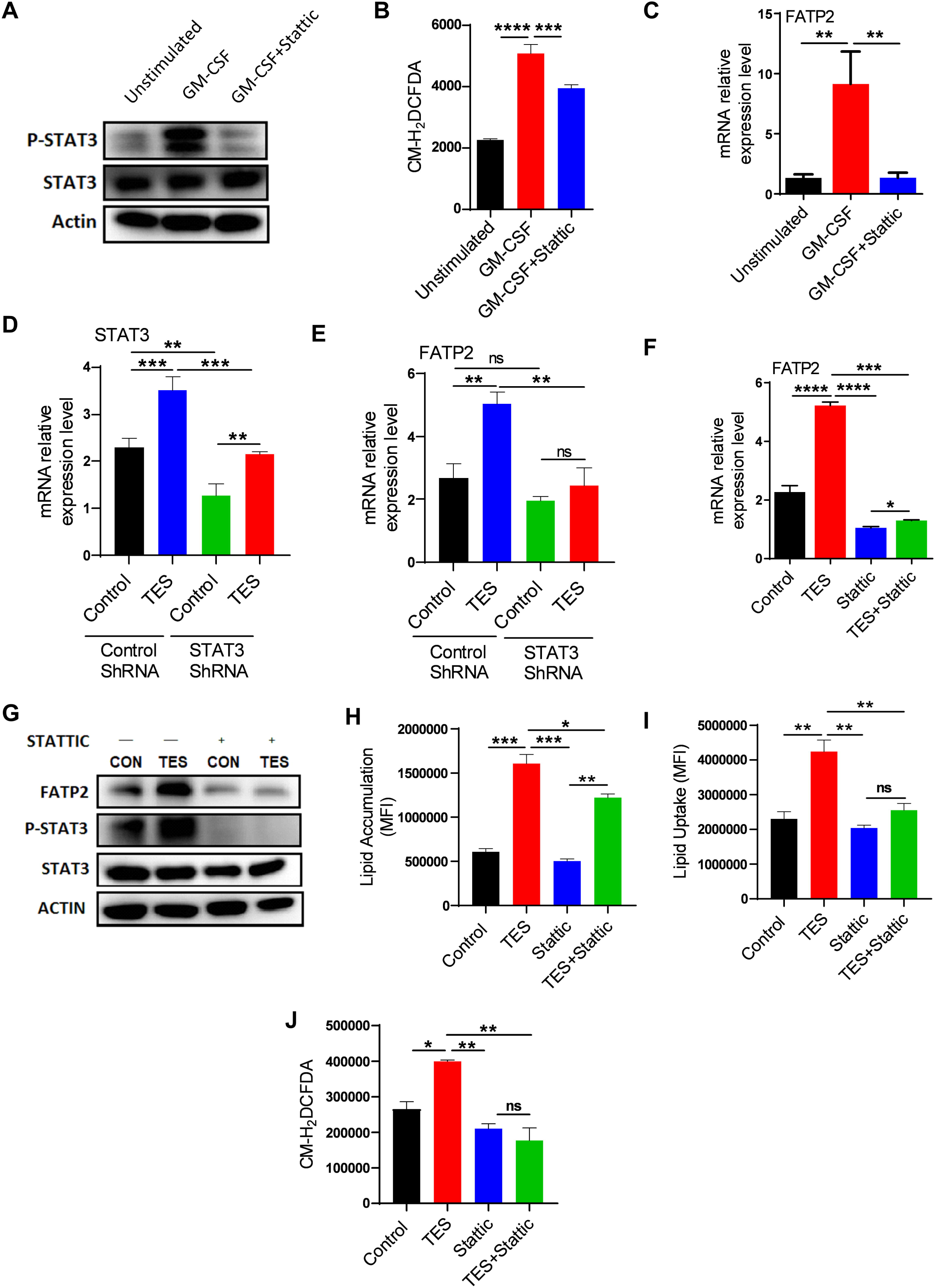
GM-CSF induces FATP2 expression by STAT3 activation in tumor MDSCs. (A-C) MDSCs derived from bone marrow cells stimulated with GM-CSF or both GM-CSF and STAT3 inhibitor, stattic (10µM) compared to unstimulated cells for 24 hours. Immunoblot analysis of STAT3 and p-STAT3 (A), Total ROS level determined by flow cytometry (B), and RT-qPCR analysis of FATP2 mRNA expression (C) in MDSCs. (D-E) mRNA expression level of STAT3 (D) and FATP2 (E) as determined by RT-qPCR in bone-marrow-derived MDSCs infected with STAT3 shRNA lentivirus. (F-J) MDSCs were generated from bone marrow cells, treated with or without TES in the presence or absence of stattic (10µM) for 24 hours. RT-qPCR analysis of FATP2 mRNA expression (F), the protein expression level of STAT3, p-STAT3, and FATP2 as determined by western blot (G), lipid accumulation (H), lipid uptake (I), and ROS generated (J) as determined by flow cytometry in MDSCs. Data are shown as representative of mean ± SEM in triplicate samples from three independent experiments. Unpaired student’s t-test or one-way ANOVA: *p < 0.05, **p < 0.01, ***p < 0.001, ****p < 0.0001, ns, no significant difference.

### 3.4 Blockade of lipid accumulation-induced ROS in MDSCs by targeting FATP2 inhibits tumor growth

To explore the potential roles of targeting FATP2 in MDSCs *in vivo*, we treated LLC and B16F10 tumor-bearing mice with lipofermata. Figures S3A-B showed the flow cytometry gating strategy for immune cells in spleens and tumors. In LLC tumor-bearing mice, lipofermata treatment significantly inhibited FATP2 mRNA and protein expression but did not change FATP4 mRNA expression in treated spleen MDSCs compared to control spleen MDSCs (Figures 4A and S4A). We then asked whether blocking FATP2 affected the distribution and function of MDSCs. We established the percentage of CD45^+^ cells from tumor tissues was not different between control and lipofermata-treated LLC tumor-bearing mice (Figure 4B). Treatment with lipofermata decreased the percentage of MDSCs and macrophages but increased DCs and CD3^+^ T cells from the tumor tissue in LLC and B16F10 tumor-bearing mice (Figures 4C and S4B). Also, lipofermata increased the population of both tumor CD4^+^ and CD8^+^ T cells gated on CD45^+^CD3^+^ cells in LLC tumor-bearing mice (Figure 4D). In the spleens of LLC and B16F10 bearing mice, lipofermata reduced the accumulation of MDSCs, PMN-MDSCs and increased the proportion of macrophages, CD3^+^ T, and NK cells while the population of M-MDSCs, DCs, and B cells remained unchanged (Figures 4E-F, and S4B-C). In addition, treatment with lipofermata increased the percentage of CD4^+^ and CD8^+^ T cells in the spleens of LLC bearing mice and CD8^+^ T cells in the spleens of B16F10 bearing mice compared to control treatment while the population of CD4^+^ T cells of B16F10 bearing mice remained unchanged (Figures 4G and S4D).

**Figure 4:**
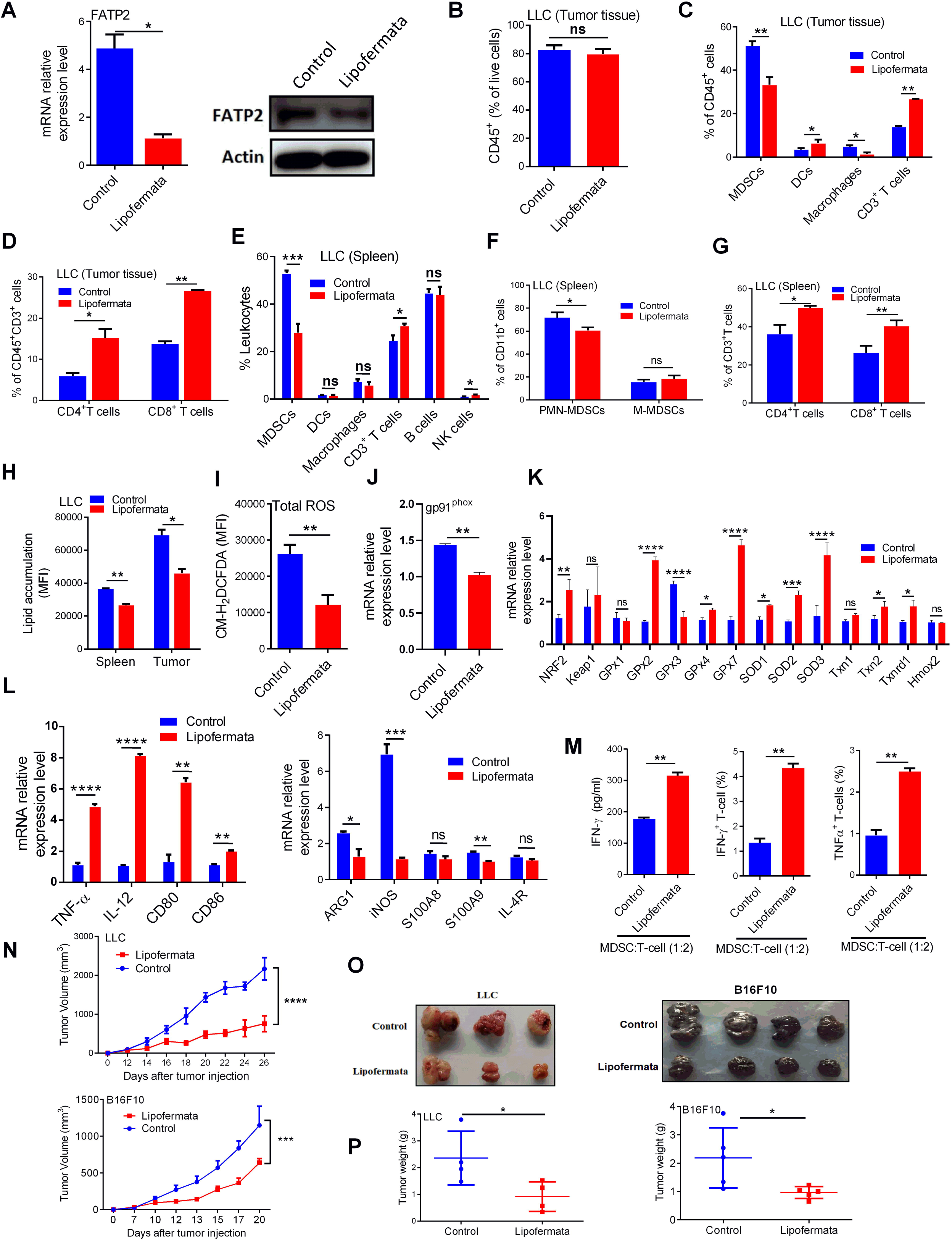
Blockade of lipid accumulation-induced ROS in MDSCs by targeting FATP2 inhibits tumor growth. (A) RT-qPCR analysis for mRNA expression (left) and immunoblot for protein expression (right) of FATP2 in spleen MDSCs isolated from control (vehicle) and lipofermata-treated LLC-bearing mice 26 days post-tumor inoculation. (B-D) Percentage of indicated leukocyte subsets among live cells in the tumor tissues of LLC tumor-bearing mice treated with vehicle (control) or lipofermata as evaluated by flow cytometry 26 days post-tumor inoculation. Leukocytes from tumor tissues were CD45^+^ cells; MDSCs were CD45^+^CD11b^+^Gr-1^+^ cells; PMN-MDSCs were CD45^+^CD11b^+^Ly6G^+^Ly6C^low^ cells; M-MDSCs were CD45^+^CD11b^+^Ly6G^−^Ly6C^high^ cells; DCs were CD45^+^CD11b^+^CD11c^+^ cells; Macrophages were CD45^+^CD11b^+^F4/80^+^ cells; CD3^+^ T cells were CD45^+^CD3^+^ cells; CD4^+^ T cells were CD45^+^CD3^+^CD4^+^CD8^−^ cells; CD8^+^ T cells were CD45^+^CD3^+^CD4^−^CD8^+^ cells. (E-G) Percentage of indicated myeloid cells and lymphocytes among live cells in the spleens of LLC tumor-bearing mice treated with vehicle (control) or lipofermata as evaluated by flow cytometry 26 days post-tumor inoculation. MDSCs were CD11b^+^Gr-1^+^ cells; PMN-MDSCs were CD11b^+^Ly6G^+^Ly6C^low^ cells; M-MDSCs were CD11b^+^Ly6G^−^Ly6C^high^ cells; DCs were CD11b^+^CD11c^+^ cells; Macrophages were CD11b^+^F4/80^+^ cells; CD3^+^ T cells were CD3^+^NK1.1^−^ cells; CD4^+^ T cells were CD3^+^CD4^+^CD8^−^ cells; CD8^+^ T cells were CD3^+^CD4^−^CD8^+^ cells; B cells were CD3^−^CD19^+^ cells; and NK cells were CD3^−^NK1.1^+^ cells. (H) MFI for lipid accumulation in MDSCs from the spleens and tumor of LLC-bearing mice treated with vehicle (control) or lipofermata. (I) Total ROS generated in spleen MDSCs from LLC tumor-bearing mice treated with control (vehicle) and lipofermata for 2 weeks. (J-K) RT-qPCR analysis of gp91^phox^ (J) and anti-oxidative genes (K) in MDSCs isolated from the spleen of LLC tumor-bearing mice treated with vehicle (control) or lipofermata (L) Gene expression profile for immune stimulatory (left) and immunosuppressive signatures (right) of MDSCs evaluated via RT-qPCR from flow cytometry sorted spleen CD11b^+^Gr1^+^ cells isolated from B16F10 tumor-bearing mice treated with control (vehicle) or lipofermata. (M) The concentration of IFN-γ released by T cells was determined via CBA (left) and percentage of IFN-γ^+^ and TNFα^+^ T cells via flow cytometry analysis (middle and right) in the co-culture of spleen MDSCs and CD3^+^ T cells at ratio 1:2, respectively. MDSCs were isolated from B16F10-bearing mice administered control (vehicle) or lipofermata for 2 weeks. CD3^+^ T cells were activated with the anti-CD3/CD28 purified antibody for 36 hours before testing. (N) Tumor growth kinetics for LLC (left) and B16F10 (right) tumor-bearing C57BL/6 mice (n=6 mice/group) treated with either control (vehicle) or lipofermata for 2 weeks. (O) Representative images for tumor tissues excised from LLC (left) and B16F10 (right) tumor-bearing C57BL/6 mice on day 26 and 20 post-tumor inoculations, respectively. (P) Tumor weight from LLC (left) and B16F10 (right) tumor-bearing mice (n=5 mice/group), respectively. Each symbol represents an individual mouse and the bar represents the mean. Data are shown as representative of mean ± SEM in triplicate samples from three independent experiments. Data represents means ± SEM of two independent experiments. Unpaired student’s t-test: *p < 0.05, **p < 0.01 ***p < 0.001, ****p < 0.0001, ns, no significant difference.

To investigate the effect of blocking FATP2 by lipofermata treatment on lipid accumulation and ROS in MDSCs from tumor-bearing mice, we incubated the spleens and tumor tissues of LLC and B16F10 bearing mice with BODIPY 493/503. Our data showed reduced lipid levels in MDSCs from both spleens and tumors of tumor-bearing mice treated with lipofermata compared to control (Figures 4H and S4E). Similarly, LC-MS analysis revealed that lipofermata reduced the accumulation of lipid molecular species such as PE, PC, PI, and TAG in B16F10 tumor-bearing mice (Figures S5A-D). Interestingly, lipofermata significantly reduced ROS production in MDSCs from the spleens of tumor-bearing mice compared to the untreated mice (Figure 4I). To confirm if lipofermata could control ROS regulatory genes in MDSCs, we determined the mRNA expression of NOX2 subunit (gp91^phox^) and antioxidant genes in MDSCs from the spleen of tumor-bearing mice compared to the control group. Similar to our in-vitro observation, lipofermata downregulated gp91^phox^ (Figure 4J), upregulated NRF2, GPx2, GPx4, GPx7, SOD1, SOD2, SOD3, Tnx2, Tnxrd1 but did not affect Keap1, GPx1, Txn1, Hmox1, Hmox2 while downregulated GPx3 mRNA expression in MDSCs from tumor-bearing mice (Figure 4K). These observations suggested that blocking FATP2 by lipofermata could suppress lipid accumulation-derived ROS in MDSCs *in vivo*.

Next, we evaluated the effect of inhibition of FATP2 by lipofermata on the suppressive activity of MDSCs in tumor-bearing mice. We analyzed the expressions of signature genes important for MDSCs function. Our data showed that MDSCs from the spleens of lipofermata-treated B16F10-bearing mice exhibited significantly decreased mRNA expression of some immunosuppressive signatures such as ARG1, iNOS, and S100A9 while IL-4R and S100A8 had no significant difference (Figure 4L). By contrast, expression of immune-stimulatory molecules such as TNF-α, IL-12, CD80, and CD86 was increased in MDSCs from B16F10-bearing mice treated with lipofermata compared to control mice (Figure 4L). The latter molecules are known to be important for T-cell activation. MDSCs from the bone marrow of lipofermata-treated B16F10-bearing mice co-cultured with naive CD3^+^ T-cells had a reduced ability to block IFN-γ or TNFα production by T-cells compared to those from control mice as determined by CBA or intracellular flow cytometry (Figures 4M and S6). The results indicate that inhibition of FATP2 expression decreased ROS-mediated immunosuppressive function in MDSCs of tumor-bearing mice but promoted T-cell activation.

To further explore the potential of pharmacological blockade of FATP2 expression in MDSCs on tumor growth, we treated LLC and B16F10-bearing mice with lipofermata and monitored the tumor kinetics. In both tumor models, treatment with lipofermata decreased tumor growth compared to control C57BL/6 mice (Figure 4N). Also, reduced tumor size and weight were noticed in the lipofermata-treated mice (Figures 4O-P). Furthermore, our observation showed that FATP2 inhibition resulted in a reduction in the spleen weight from lipofermata treated LLC-bearing mice compared to untreated mice but no change in the spleen weight from lipofermata treated B16F10-bearing mice compared to untreated mice (Figures S7A-B). Surprisingly, lipofermata failed to reduce B16F10 tumor growth in immunocompromised balb/c nude mice (Figure S7C). Similarly, inhibiting FATP2 in LLC and B16F10 cells did not alter their proliferation *in-vitro* (Figures S7D-E). On the contrary, purified MDSCs from the bone marrows of B16F10-bearing mice demonstrated a decrease in cell proliferation on treatment with lipofermata in a dose-dependent manner (Figure S7F). These results indicate that blockade of lipid accumulation**-**induced ROS in MDSCs by targeting FATP2 inhibits tumor growth.

### 3.5 Pharmaceutical blockade of FATP2 in MDSCs enhances anti-PD-L1 tumor immunotherapy

Next, we explored whether lipofermata could demonstrate a synergistic effect with the anti-PD-L1 antibody in delaying tumor progression. C57BL/6 mice were injected with B16F10 or LLC cells and allowed to grow until the tumor was palpable on days 6 and 12 respectively. Mice received lipofermata daily while anti-PD-L1 antibody was administered every three days, starting from day 6 and 12 following injection of B16F10 and LLC respectively. Tumor growth kinetics revealed that either lipofermata or anti-PD-L1 antibody alone had an anti-tumor effect compared to the control (Figure 5A). Interestingly, the combination of anti-PD-L1 and lipofermata drastically delayed tumor progression compared to the monotherapy (Figure 5A). The representative images on day 20 and 28 post tumor inoculation for B16F10 and LLC respectively showed a significant reduction in the tumor sizes from combination-treated mice (Figure 5B) corroborated with decreased tumor weights (Figure 5C) compared to the control and single therapy. Also, there was no difference in the body weight of all mice receiving different treatments compared to the control (Figure 5D). Moreover, treatment did not result in splenomegaly in B16F10-bearing mice, rather the combination treatment resulted in more reduced spleen weight compared to the control and single agents (Figure 5E). Although spleen weight was not recorded for LLC-bearing mice. Therefore, our data suggest that pharmaceutical blockade of FATP2 in MDSCs enhanced anti-PD-L1 tumor immunotherapy.

**Figure 5:**
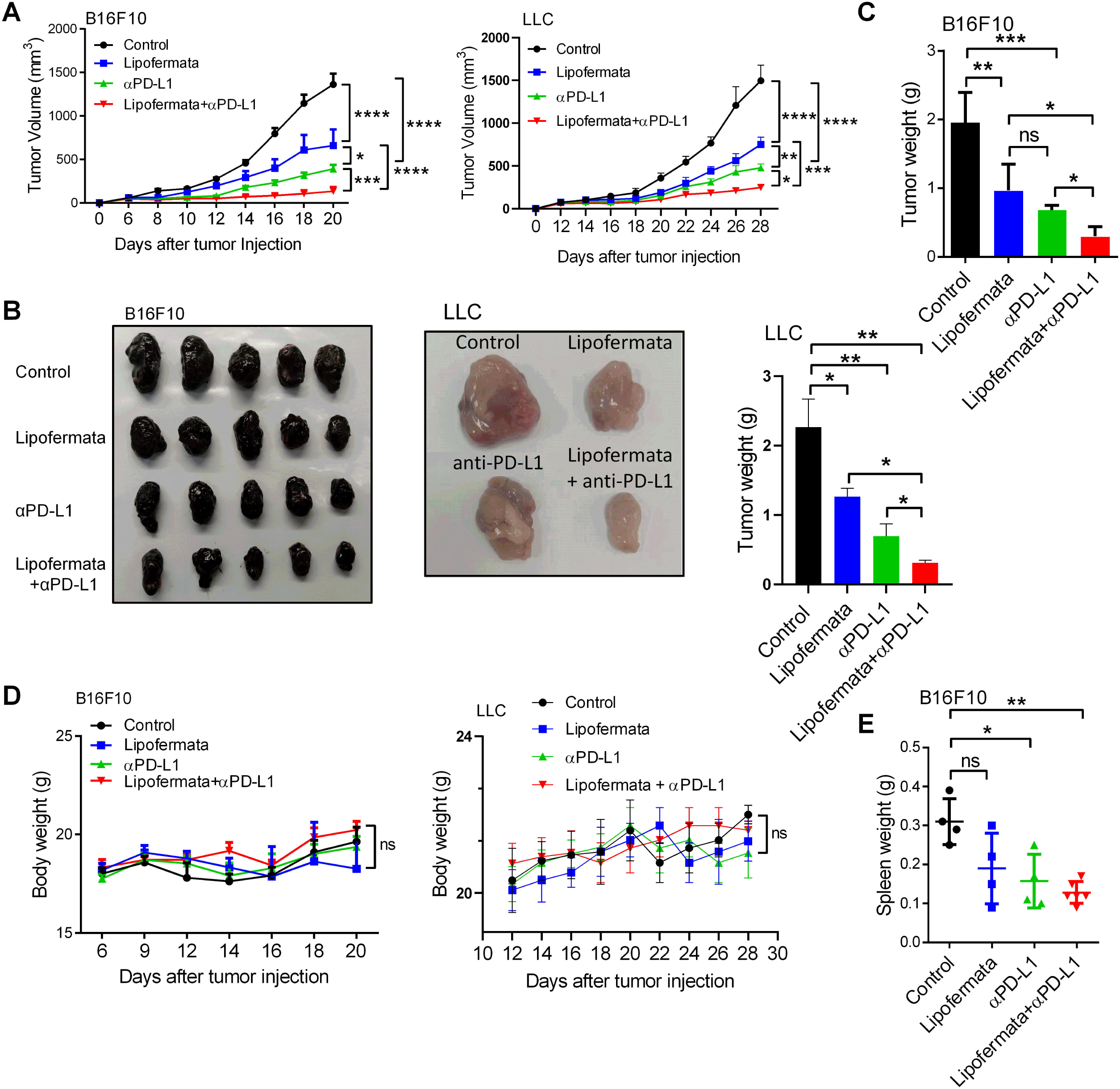
Pharmaceutical blockade of FATP2 in MDSCs enhances anti-PD-L1 immunotherapy. (A) Tumor growth curve of B16F10 and LLC tumor-bearing mice (n=7 mice/group) treated with lipofermata daily, anti-PD-L1 antibody every three days starting from day 6 and 12 respectively, or anti-PD-L1 antibody combined with lipofermata. Results are means ± SEM pooled from two independent experiments. (B) Representative images of tumor tissues excised from lipofermata, anti-PD-L1 antibody, and the combination-treated B16F10 and LLC tumor-bearing mice (n=5 mice/group). (C) Tumor weight recorded from B16F10 and LLC tumor-bearing mice (n=5 mice/group) that received either lipofermata daily, anti-PD-L1 antibody, or anti-PD-L1 antibody combined with lipofermata therapy on day 20 and 28 after B16F10 and LLC inoculation respectively. (D) Body weight kinetics of B16F10 and LLC tumor-bearing mice (n=7 mice/group) treated with either lipofermata daily, anti-PD-L1 antibody every three days post-tumor inoculation, or anti-PD-L1 antibody combined with lipofermata therapy. (E) Spleen weight from B16F10 tumor-bearing mice (n=4 mice/group) treated with either lipofermata daily, anti-PD-L1 antibody, or anti-PD-L1 antibody combined with lipofermata therapy on day 20 post-tumor inoculation. Data represents means ± SEM of two independent experiments. Unpaired student’s t-test, one-way ANOVA or two-way ANOVA: *p < 0.05, **p < 0.01 ***p < 0.001, ****p < 0.0001, ns, no significant difference.

### 3.6 FATP2 blockade augments anti-PD-L1 tumor immunotherapy through the activation of T-cells and inhibition of MDSCs suppressive role

We moved on to investigate how treatment with FATP2 inhibitor enhanced the efficacy of PD-L1 checkpoint blockade in tumor-bearing mice. Considering the anti-tumor potential of T-cells, we evaluate the percentage of tumor-infiltrating CD8^+^ and CD4^+^ T-cells in LLC-bearing mice treated with either anti-PD-L1 antibody or lipofermata alone, or both anti-PD-L1 antibody and lipofermata. Our data showed that either lipofermata or anti-PD-L1 antibody alone increased the population of CD8^+^ and CD4^+^ T-cells compared to the control group (Figure 6A); importantly, the combination of lipofermata and anti-PD-L1 treatment significantly increased the proportion of CD8^+^ and CD4^+^ T-cells compared to the monotherapy (Figure 6A). Next, we asked if T-cells activation markers such as CD107a and CD69 known to be involved in anti-tumor activity is upregulated in the combined therapy group. Flow cytometry data showed that both lipofermata and anti-PD-L1 antibody increased the cell surface expression of CD107a but only lipofermata increased the surface expression of CD69 on tumor-infiltrating CD8^+^ and CD4^+^ T-cells while anti-PD-L1 treatment did not affect CD69 expression compared to the control group (Figures 6B-C). Interestingly, lipofermata enhanced PD-L1 blockade-mediated therapy via significantly increased expression of CD107a on CD8^+^ and CD4^+^ T-cells compared to either treatment alone (Figure 6B). On the contrary, combined therapy of lipofermata and anti-PD-L1 antibody did not demonstrate a synergistic effect on the expression of CD69 on T-cells (Figure 6C). Furthermore, we decided to find out if lipofermata could regulate PD-L1 expression on immune cells thereby augments PD-L1 blocked-mediated therapy. In tumor-infiltrating CD45^+^ cells, either lipofermata or anti-PD-L1 antibody reduced the surface expression of PD-L1 compared to the control (Figure 6D) while the combination of anti-PD-L1 and lipofermata treatment demonstrated a synergistic effect to further reduce PD-L1 expression (Figure 6D). Importantly, our data showed that either lipofermata or anti-PD-L1 antibody decreased surface expression of PD-L1 on CD8^+^ T-cells compared to the control while combination treatment further reduced PD-L1 expression compared to monotherapy. (Figure 6E). Although anti-PD-L1 treatment slightly reduced PD-L1 surface expression on tumor-infiltrating CD4^+^ T-cells, lipofermata increased its expression compared to control while the combined therapy had no significant difference compared to the lipofermata-treated group alone (Figure 6E). Altogether, these suggest the activation of tumor-infiltrating CD8^+^ T-cells in mice treated with a combination of lipofermata and anti-PD-L1 antibody for enhanced cytotoxic activity thus delaying tumor growth.

**Figure 6:**
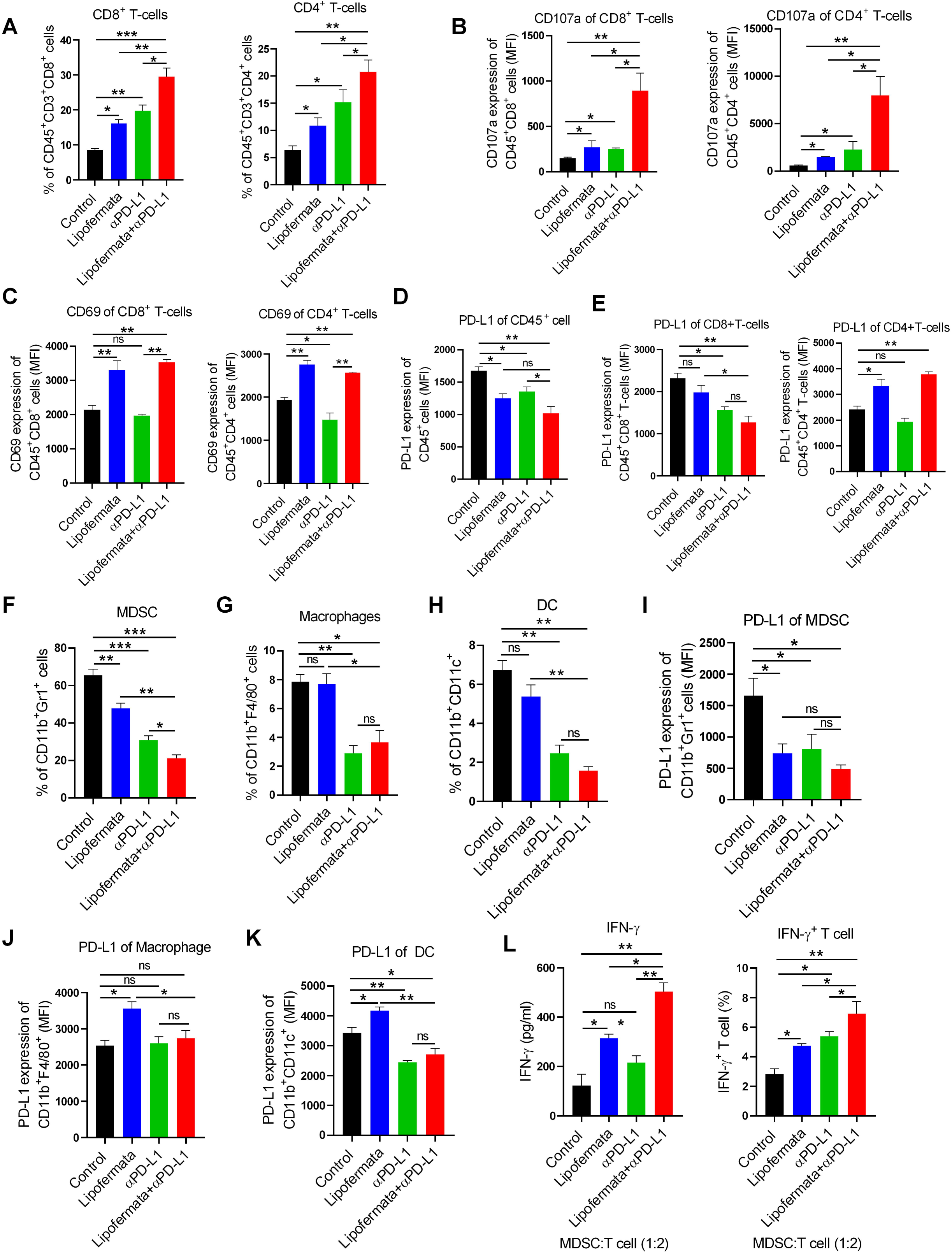
FATP2 blockade augments anti-PD-L1 tumor immunotherapy through the activation of T-cells and inhibition of MDSCs suppressive role. (A) Percentage of CD45^+^CD3^+^CD8^+^ and CD45^+^CD3^+^CD4^+^ T cells present in leukocytes live cells from tumor tissue of LLC-bearing mice treated with or without lipofermata, anti-PD-L1 antibody (clone 10F.9G2, Bio X Cell), or combined therapy evaluated via flow cytometry. (B-C) MFI for CD107a (A) and CD69 (B) expression on CD45^+^CD8^+^ and CD45^+^CD4^+^ T cells in tumors of LLC-bearing mice treated with or without lipofermata, anti-PD-L1, or combined therapy. (D-E) MFI for surface expression of PD-L1 (clone MIH5, Invitrogen) on tumor-infiltrating CD45^+^ cells (D) and CD45^+^CD8^+^ or CD45^+^CD4^+^ T cells (E) from LLC-bearing mice treated with or without mono- or combination therapy. (F-H) Proportion of myeloid cells present in the spleen of LLC-bearing mice treated with or without lipofermata, anti-PD-L1, or combined therapy evaluated on flow cytometry on day 28 post-tumor injection. MDSCs were CD11b^+^Gr-1^+^ cells; Macrophages were CD11b^+^F4/80^+^ cells and DCs were CD11b^+^CD11c^+^ cells. (I-K) MFI for surface expression of PD-L1 present in MDSCs (I), macrophages (J), and DCs (K) in the spleen of LLC-bearing mice receiving mono- or combination treatment. (L) The concentration of IFN-γ released by T-cells was determined via CBA (left) and percentage of IFN-γ^+^ T-cells via flow cytometry analysis (right) in the co-culture of spleen MDSCs and CD3^+^ T-cells at ratio 1:2, respectively. MDSCs were isolated from LLC-bearing mice administered control (vehicle), lipofermata, anti-PD-L1, or combination therapy and sacrificed on day 28 post-tumor injection. CD3^+^ T-cells were activated with the anti-CD3/CD28 purified antibody for 36 hours before testing. Data are shown as representative of mean ± SEM in triplicate samples from three independent experiments. Data represents means ± SEM of two independent experiments. Unpaired student’s t-test: *p < 0.05, **p < 0.01 ***p < 0.001, ****p < 0.0001, ns, no significant difference.

Next, we explored how the combined therapy will affect suppressive myeloid cell populations and their function on T-cells anti-tumor activity. Our data showed a reduced proportion of MDSCs, macrophages, and DCs from the spleens of LLC-bearing mice treated with anti-PD-L1 antibody compared to the control (Figure 6F-H). However, lipofermata-treatment significantly reduced MDSCs population but had no significant effect on macrophages and DCs compared to the control (Figure 6F-H). Interestingly, combination therapy substantially decreased MDSCs population and slightly reduced DCs proportion but not macrophages compared to anti-PD-L1 therapy alone (Figure 6F-H). Furthermore, anti-PD-L1 blockade therapy reduced PD-L1 surface expression on MDSCs and DCs but did not affect that of macrophages compared to the control (Figure I-K). Notably, lipofermata inhibited the expression of PD-L1 on MDSCs (Figure 6I), unlike macrophages and DCs which had increased expression (Figure 6J-K). In the combination treatment, MDSCs showed decreased surface expression of PD-L1 although this was not significant compared to the anti-PD-L1 treated mice alone (Figure 6I). On the contrary, data show that the expression of PD-L1 on macrophages and DCs in the combined therapy group were not different from that of mice which received anti-PD-L1 antibody alone (Figures 6J-K). MDSCs from the spleen of either lipofermata or anti-PD-L1 treated LLC-bearing mice co-cultured with naïve CD3^+^ T cells had decreased ability to block IFN-γ production by T-cells compared to those from untreated mice (Figure 6L). More importantly, MDSCs from mice treated with the combination of anti-PD-L1 antibody and lipofermata demonstrated a synergistic effect for reduced capability to hinder IFN-γ production by T-cells compared to the monotherapy (Figure 6L). Collectively, our data show that tumor cells-derived GM-CSF activated STAT3 signaling, leading to upregulation of FATP2 expression, thus an increase in tumor-derived lipid uptake and accumulation in MDSC. Consequently, this FATP2-induced lipid accumulation enhanced mitochondrial function and activated ROS thereby promoting the immunosuppressive function of MDSCs on T-cells, thus resulting in tumor progression. However, pharmaceutical blockade of FATP2 in MDSCs enhanced anti-PD-L1 tumor immunotherapy via the activation of T-cells for enhanced production of IFN-γ (Figure 7).

**Figure 7:**
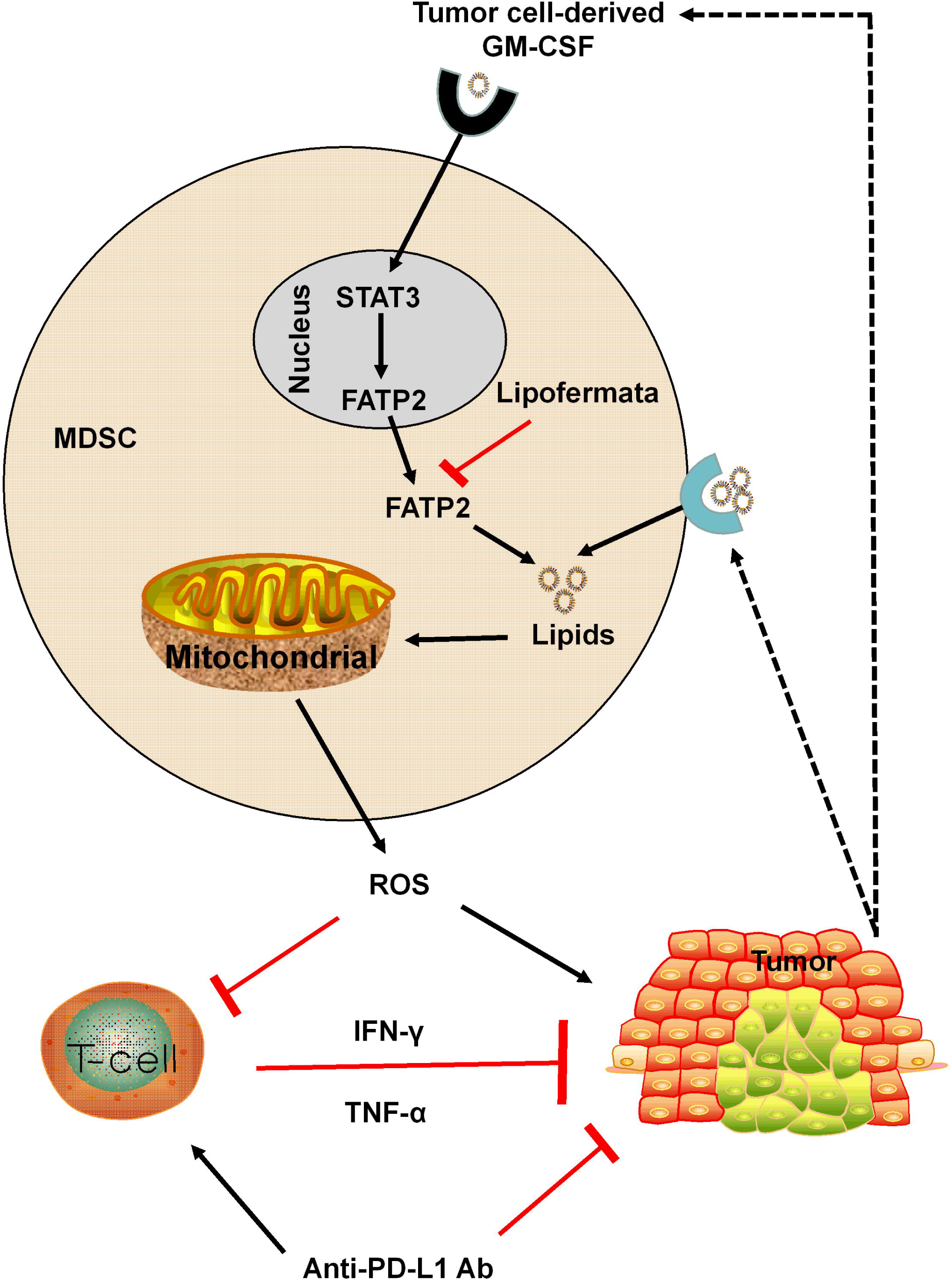
Proposed mechanism of action for this study. MDSC – Myeloid-derived suppressor cells; STAT3 - signal transducer and activator of transcription 3; FATP2 – Fatty acid transport protein 2; GM-CSF – Granulocyte-macrophage colony-stimulating factor; ROS – Reactive oxygen species; TNFα – Tumor necrosis factor-alpha; IFN-γ – Interferon-gamma; anti-PD-L1 Ab – Anti-programmed death-ligand 1 antibody. The black arrow denotes activation while the red arrow represents inhibition.

## 4. Discussion

Here we report that FATP2 regulates MDSCs immunosuppressive function via controlling ROS production and differentiation of MDSCs in tumor-bearing mice. Pharmaceutical blockade of FATP2 expression in MDSCs by lipofermata decrease lipid accumulation and ROS while upregulating the expression of antioxidant genes, promote differentiation of MDSCs to immune-stimulatory cells thus result in reduced MDSCs immunosuppressive activity and consequent inhibition of tumor growth in mice. Moreover, this study shows for the first time that inhibiting FATP2 in MDSCs further enhances anti-PD-L1 tumor immunotherapy through increased T-cells population, upregulated expression of CD107a, and reduced PD-L1 expression on tumor-infiltrating CD8^+^ T-cells. Furthermore, combined therapy of FATP2 inhibition and anti-PD-L1 antibody decreased the immunosuppressive function of MDSCs on CD3^+^ T-cells rather enhances T-cell’s ability to release IFN-γ for anti-tumor effect.

Previous studies established the involvement of lipid-induced ROS in controlling immune cell polarization and activities [9, 36–38]. Tumor-derived MDSCs prefer lipid metabolism as their energy source [9, 20]. Increased ROS drives MDSCs immunosuppressive activity on T-cells as well as inhibit its differentiation to other myeloid cells [39]. Elevated level of ROS is a major characteristic of MDSCs present in tumor-bearing mice and cancer patients [40–42]. The immune-suppressive function of MDSCs in tumor-bearing mice and cancer patients can be overcome by blocking ROS production in MDSCs [43]. However, it remains unknown if regulating FATP2 expression of MDSCs could suppress ROS production in MDSCs. We report here that unsaturated fatty acid and TES induced ROS production (Figures 1A-E) while targeting FATP2 decrease ROS in tumor-induced MDSCs (Figure 1P). Though ROS can be generated in cells via several mechanisms, the major source of ROS in leukocytes including MDSCs is NADPH oxidase (NOX2) [44]. Here we show that lipofermata repressed gp91^phox^ expression (a subunit of NOX2) in TES-induced MDSCs as well as spleen MDSCs from tumor-bearing mice (Figures 1Q and 4J). This corroborates previous work that MDSCs from gp91^phox^ deficient tumor-bearing mice demonstrated reduced ROS level, failed to suppress T-cell function, differentiated to F4/80^+^Gr1^-^ macrophages and CD11b^+^CD11c^+^ DCs, and slightly inhibited tumor growth[44]. From our study, this suggests decreased gp91^phox^ expression in MDSCs by FATP2 inhibition to be crucial for ROS-mediated function of MDSCs. Also, targeting FATP2 expression in TES-induced MDSCs drastically upregulated the expression of ROS regulatory genes as shown in Figure 1R. Similar to our in-vitro observation, tumor-bearing mice treated with lipofermata show decreased MDSCs ROS production with concomitant increased expression of antioxidant genes (Figures 4I and 4K). NRF2 is a master regulator of antioxidant genes and MDSCs isolated from NRF2-deficient mice had been reported to demonstrate elevated ROS level thereby enhanced tumor growth [45, 46]. Importantly, in this study lipofermata upregulated the expression of NRF2 both in-vitro and in-vivo (Figures 1R and 4K). These findings reveal a new role of FATP2 in regulating ROS-mediated immunosuppressive function of MDSCs through NRF2 induction of anti-oxidative genes.

In addition, MDSCs mediate their suppressive effect on T-cells via the generation of nitric oxide by enhanced ARG1 and iNOS expression [47]. MDSCs overexpression of ARG1 and iNOS promotes the metabolism of arginine thereby depleting arginine required for T-cells anti-tumor response [48, 49]. We showed that FATP2 inhibition in tumor-bearing mice repressed the expression of ARG1 and iNOS in spleen MDSCs (Figure 4L). This is contrary to the observation of Veglia et al., who reported no significant difference in the expression of ARG1 and iNOS in spleen PMN-MDSCs from FATP2 knock out (KO) EL4 tumor-bearing mice [21]. This could be dependent on the tumor model from which MDSCs were isolated. Consequently, our study highlights targeting FATP2 expression in MDSCs downregulated immunosuppressive signatures thereby decreased their suppressive role on T-cells and enhanced the anti-tumor effect of T-cells.

GM-CSF is a key modulator of MDSCs generation and activation [32, 50]. Nonetheless, it’s been reported that *in-vitro* generation of functional MDSCs from naive progenitor cells requires a minimum of two signal mediators [51, 52]. Our study shows tumor cells derived-GM-CSF promote lipid accumulation as well as fatty acid uptake in MDSCs. On the contrary, IL-6, an immunosuppressive cytokine expressed in MDSCs, slightly increased lipid accumulation while it did not affect fatty-acid uptake in MDSCs. More so, combination of GM-CSF and IL-6-induced MDSCs failed to show a synergistic effect in their ability for fatty acid uptake or lipid accumulation (Figures 2F-G). Although unsaturated fatty acid slightly induced IL-6 expression in BM-derived MDSCs to promote the immunosuppressive role of MDSCs (Figures S2B-C). Furthermore, our data show that GM-CSF is mainly derived from tumor microenvironment confirming exogenous fatty acid mediating lipid accumulation in MDSCs (Figures 2A-C). On the other hand, IL-6 is majorly released by MDSCs suggesting that endogenous fatty acid synthesis could partly contribute to accumulated lipid in MDSCs observed following stimulation with IL-6 (Figures 2A-B and 2D-E). This indicates the important role of IL-6 in driving lipid accumulation in MDSCs and requires more studies to elucidate the mechanism.

Moreover, GM-CSF is a critical cytokine for regulating FATP2 expression in MDSCs. GM-CSF regulates the expression of FATP2 through the activation of STAT5 signaling pathway in PMN-MDSCs [21]. However, our study shows that tumor cells-derived GM-CSF triggered FATP2 expression via activation of STAT3 signaling pathway, thereby promoting lipid accumulation-induced ROS in MDSCs. Previous studies showed GM-CSF/JAK2/STAT3 signaling axis mediates MDSCs immunosuppression in tumor metastasis and blockade of this pathway enhanced antitumor activity [34, 35]. Also, our group earlier reported the ability of polyunsaturated fatty acid to promote MDSCs expansion via activation of STAT3 signaling pathway and targeting STAT3 decreased MDSCs accumulation and their suppressive function [21, 53]. More importantly, we show here that blockade of GM-CSF signal or STAT3 activation downregulates FATP2 expression, thus resulting in reduced lipid accumulation in MDSCs. Besides, inhibiting STAT3 expression in MDSCs decreased its ability for lipid uptake and consequently blocked ROS generation in TES-induced MDSCs.

Previous study showed that FATP2 is absent in DC, macrophages, M-MDSCs but slightly present in CD8^+^ T cells and abundantly expressed in PMN-MDSCs from EL-4 tumor-bearing mice [21]. On the contrary, we report that using lipofermata to specifically target FATP2 in LLC tumor-bearing mice had a direct or indirect effect on several tumor-infiltrating immune cells (Figures 4C-D). This led to reduced accumulation of MDSCs and macrophages while increasing the proportion of DCs and T cells in tumor-infiltrating CD45^+^ cells in our study compared to an earlier work that reported reduced PMN-MDSCs alone in FATP2 KO mice [21]. Here lipofermata demonstrates the ability to reduce the percentage of macrophages, another immunosuppressive phenotype in tumor milieu which underlines its potential as a promising anti-immune inhibitory agent (Figure 4C). This corroborates a recent report that regulating lipid droplet in TAM by targeting diacylglycerol transferases (DGAT), an enzyme needed for the synthesis of triglycerides, inhibited tumor growth in mice [54]. Our gene expression analysis shows that FATP2 blockade in MDSCs isolated from spleens of lipofermata-treated tumor-bearing mice upregulated the expression of IL-12, CD80 and CD86 (Figure 4K). These proteins are co-stimulatory molecules expressed on antigen-presenting cells (APCs) and known for their role in T cells activation [55, 56]. Also, IL-12 is a potent inducer of IFN-γ production by T and NK cells for cytotoxic activity [57]. Altogether, our results suggest an unreported role of lipofermata in decreasing macrophages population and promoting the differentiation of MDSCs from its immunosuppressive phenotype to APCs such as DCs.

More importantly, the combination of lipofermata and anti-PD-L1 treatment led to decreased MDSCs population and surface expression of PD-L1 in spleens from LLC-bearing mice (Figures 6F and 6I). This almost completely abrogated MDSCs immunosuppressive effect on T-cells function (Figure 6L). Besides, blockade of FATP2 in MDSCs by lipofermata treatment in tumor-bearing mice enhances anti-PD-L1 tumor immunotherapy via activation of tumor-infiltrating CD8^+^ T-cells through increased expression of CD107a, enhanced ability for IFN-γ production, and reduced PD-L1 surface expression (Figures 6A-B, 6E, and 6L). This led to delay in LLC progression compared to monotherapy (Figures 5A-C).

## 5. Conclusion

We have discovered that FATP2 controls MDSCs immunosuppressive function in part through regulating ROS production and MDSCs differentiation. The potential of FATP2 in modulating lipid-derived ROS is crucial for tumorigenesis since the pharmaceutical blockade of FATP2 in MDSCs by lipofermata leads to a significant reduction in mouse tumor progression. More importantly, our findings highlight the possibility of therapeutic inhibition of FATP2 in MDSCs by lipofermata in cancer patients to overcome obstacles for effective anti-PD-L1 cancer immunotherapy.

## CRediT authorship contribution statement

**Dehong Yan**: Conceptualization; **Adeleye O. Adeshakin, Funmilayo O. Adeshakin** and **Wan Liu**: Data curation; **Adeleye O. Adeshakin** and **Wan Liu**: Formal analysis; **Adeleye O. Adeshakin, Funmilayo O. Adeshakin** and **Wan Liu**: Investigation; **Funmilayo O.Adeshakin and Menqgi Zhang**: Methodology; **Adeleye O. Adeshakin and Guizhong Zhang**: Software; **Adeleye O. Adeshakin, Lukman O. Afolabi and Menqgi Zhang**: Validation; **Adeleye O. Adeshakin, Funmilayo O. Adeshakin, Lukman O. Afolabi, Lulu Wang, Zhihuan Li, and Lilong Lin**: Visualization; **Adeleye O. Adeshakin**: Roles/Writing - original draft; **Xiaochun Wan and Dehong Yan**: Funding acquisition; Project administration; Resources; Supervision; Writing - review & editing.

## Acknowledgments

Adeleye Oluwatosin Adeshakin is sponsored by the University of Chinese Academy of Sciences (UCAS) and Shenzhen Institute of Advanced Technology scholarship for international students. We thank members of the Public Technology Service Cores at the Shenzhen Institutes of Advanced Technology, Chinese Academy of Sciences, for their technical assistance. We also appreciate the support from Zhan Wugen of the Shenzhen Institutes of Advanced Technology, Chinese Academy of Sciences, for flow cytometry cell sorting.

## Supplementary data

**Figure S1:**
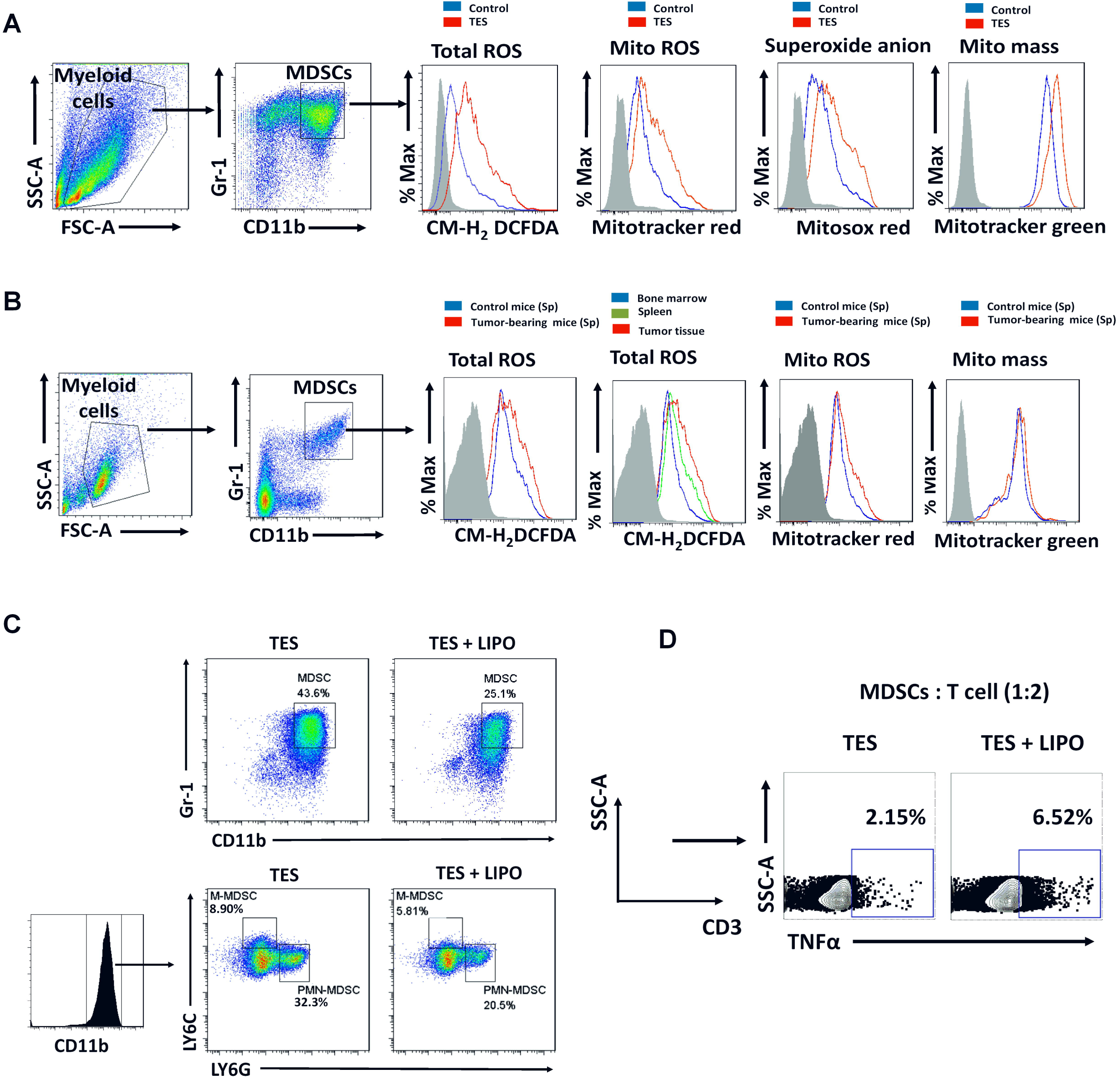
FATP2 regulates ROS and immunosuppressive function of MDSCs. (A) Representative histograms for flow cytometry analysis of total ROS, mitochondrial ROS, superoxide anion, and mitochondrial mass in MDSCs from GM-CSF stimulated bone marrow cells cultured in the presence or absence of TES for 24 hours as illustrated in Figures 1A-D. (B) Representative histograms for flow cytometry analysis of total ROS, mitochondrial ROS, and mitochondrial mass respectively in spleen CD11b^+^Gr-1^+^ cells of tumor-free mice and spleen MDSCs of B16F10 tumor-bearing mice or total ROS in the spleen, bone marrow, and tumor MDSCs of B16F10 tumor-bearing mice as illustrated in figures 1G-I. (C) Flow cytometry gating strategy for total MDSCs, PMN-MDSCs, and M-MDSCs from GM-CSF-induced bone marrow cells treated TES with or without lipofermata for 48 hours as illustrated in figure 1O. (D) Flow cytometry gating strategy for TNFα^+^ T cells in co-culture of MDSCs and CD3^+^ T cells at ratio 1:2 respectively as illustrated in Figure 1T (right). MDSCs from GM-CSF-induced bone marrow cells treated TES with or without lipofermata for 48 hours. CD3^+^ T cells were activated with the anti-CD3/CD28 purified antibody for 36 hours before testing. MDSCs were CD11b^+^Gr-1^+^ cells; PMN-MDSCs were CD11b^+^Ly6G^+^Ly6C^low^ cells; M-MDSCs were CD11b^+^Ly6G^−^Ly6C^high^ cells; Tumor explanted supernatant-TES; Linoleic acid-LA; Oleic acid–OA; α-Linolenic acid–ALA; Stearic acid–SA; Mitochondrial-Mito, Lipofermata-LIPO.

**Figure S2:**
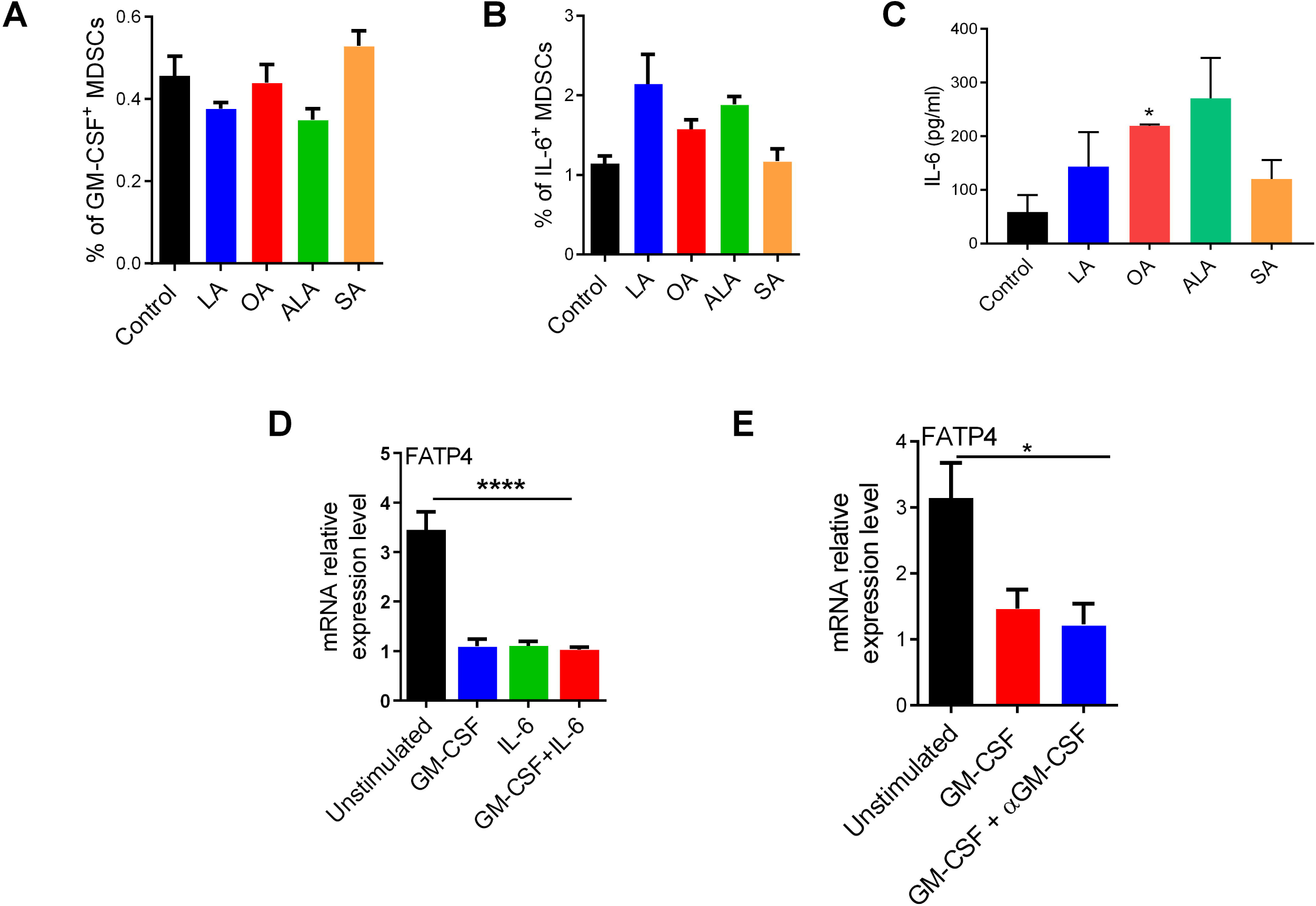
GM-CSF or IL-6 does not trigger FATP4 expression in MDSCs. (A-C) Flow cytometry analysis showing the percentage of GM-CSF (A), IL-6 (B), and IL-6 release as determined by CBA (C) in bone-marrow-derived MDSC cultured in the presence of indicated fatty acids for 24 hours. (D) RT-qPCR analysis for FATP4 mRNA expression of bone marrow-derived MDSCs stimulated with GM-CSF, IL-6, or both cytokines. (E) RT-qPCR analysis for FATP4 mRNA expression of bone marrow-derived MDSCs stimulated in the presence or absence of GM-CSF and blocked with or without anti-GM-CSF antibody (5µg/ml). Linoleic acid-LA; Oleic acid–OA; α-Linolenic acid–ALA; Stearic acid–SA; Data are shown as representative of mean ± SEM in triplicate samples from three independent experiments. Unpaired student’s t-test or one-way ANOVA: *p < 0.05, ****p < 0.0001.

**Figure S3:**
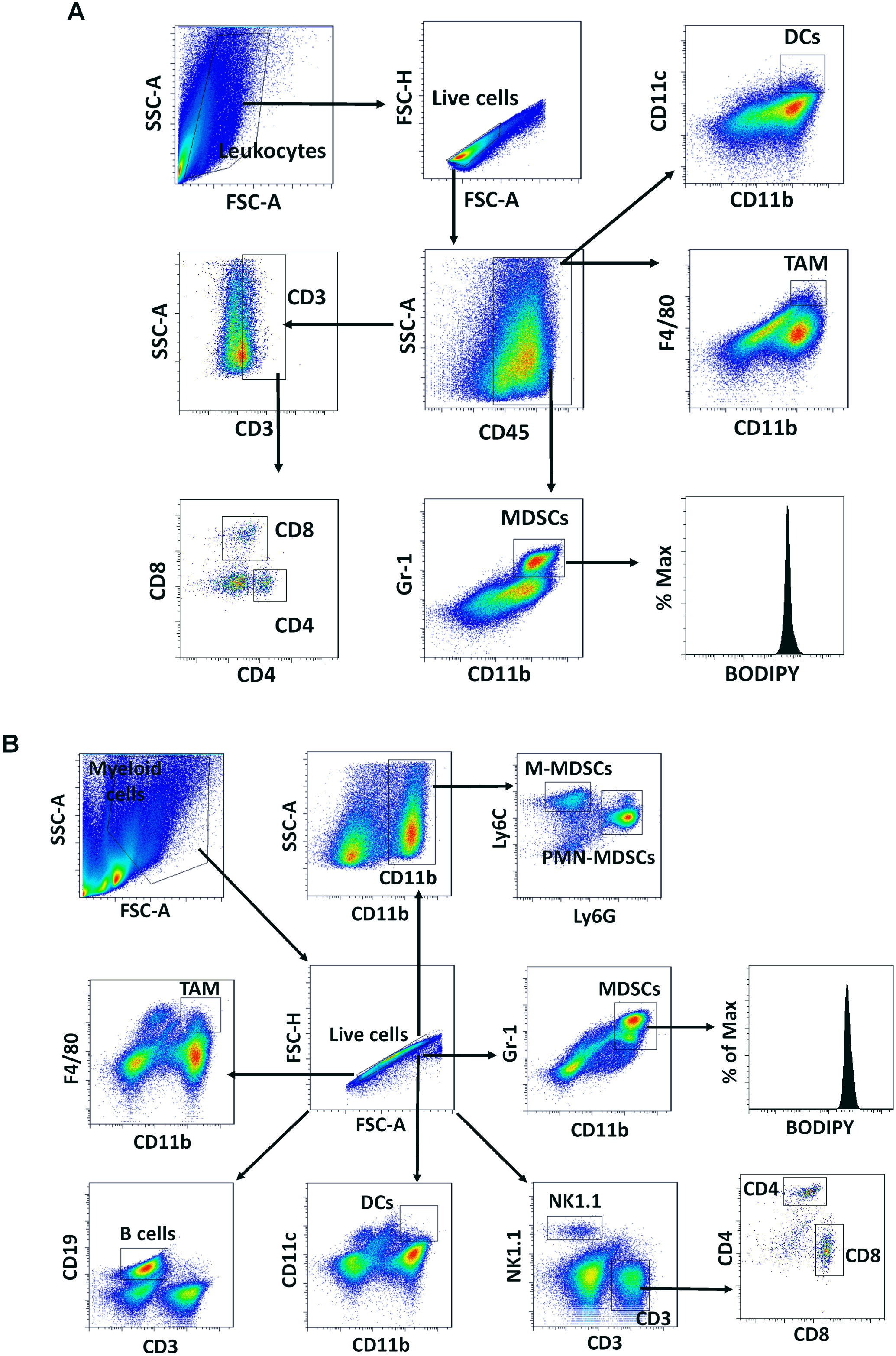
Flow cytometry plots illustrating the gating strategy used for leukocytes analysis. (A) Representative flow cytometry plots showing gating strategy for tumor-infiltrating leukocytes in Figures 4B-D and 4H. (B) Representative flow cytometry plots illustrating the gating strategy for splenocytes in Figures 4E-H. Leukocytes from tumor tissues were CD45^+^ cells: MDSCs were CD45^+^CD11b^+^Gr-1^+^ cells; PMN-MDSCs were CD45^+^CD11b^+^Ly6G^+^Ly6C^low^ cells; M-MDSCs were CD45^+^CD11b^+^Ly6G^−^Ly6C^high^ cells; DCs were CD45^+^CD11b^+^CD11c^+^ cells; macrophages were CD45^+^CD11b^+^F4/80^+^ cells; CD3^+^ T cells were CD45^+^CD3^+^ cells; CD4^+^ T cells were CD45^+^CD3^+^CD4^+^CD8^−^ cells; CD8^+^ T cells were CD45^+^CD3^+^CD4^−^CD8^+^ cells; For splenocytes: MDSCs were CD11b^+^Gr-1^+^ cells; G-MDSCs were CD11b^+^Ly6G^+^Ly6C^low^ cells; M-MDSCs were CD11b^+^Ly6G^−^Ly6C^high^ cells; DCs were CD11b^+^CD11c^+^ cells; macrophages were CD11b^+^F4/80^+^ cells; CD3^+^ T cells were CD3^+^NK1.1^−^ cells; CD4^+^ T cells were CD3^+^CD4^+^CD8^−^ cells; CD8^+^ T cells were CD3^+^CD4^−^CD8^+^ cells; B cells were CD3^−^CD19^+^ cells; and NK cells were CD3^−^NK1.1^+^ cells.

**Figure S4:**
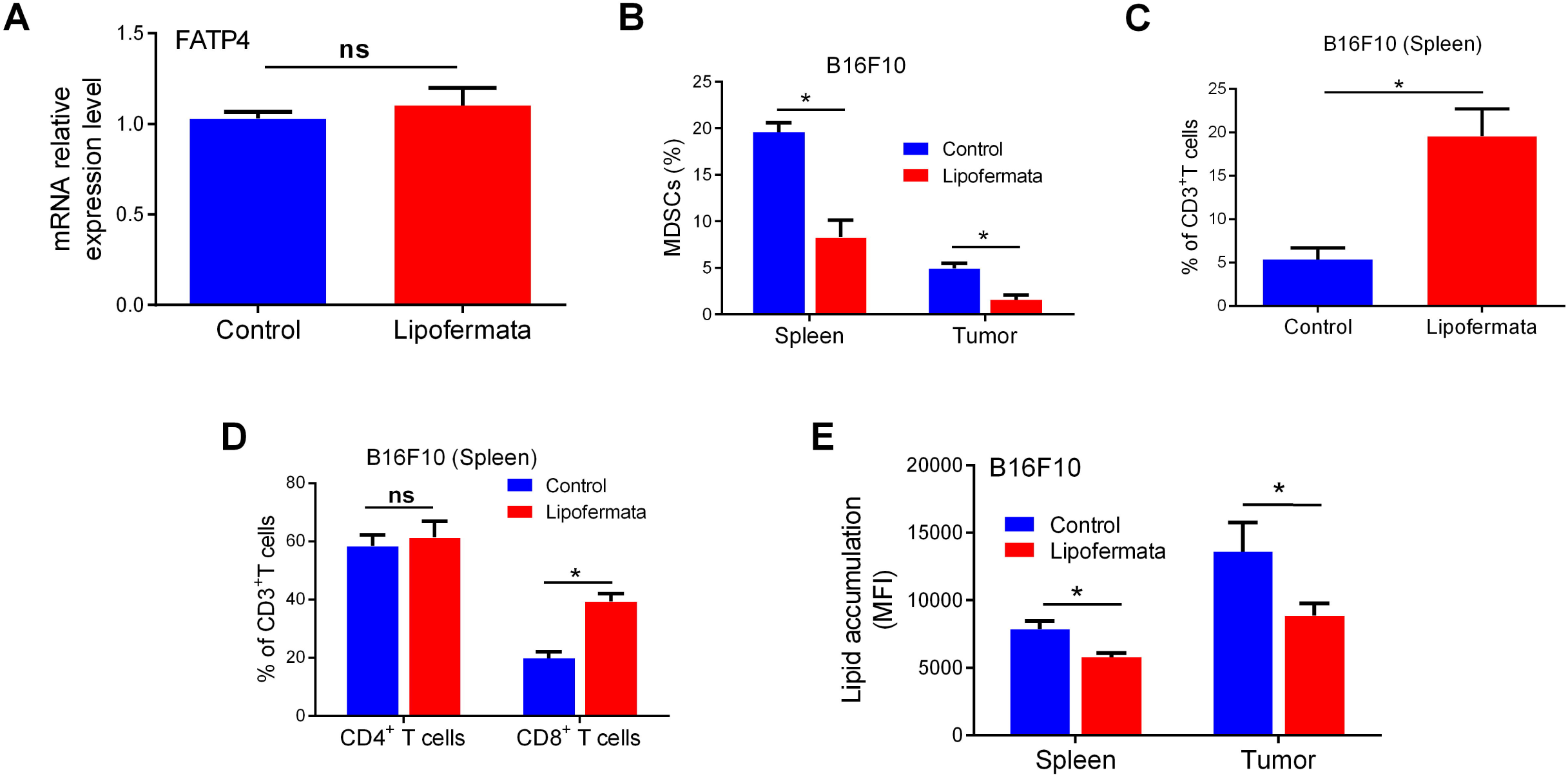
Impact of lipofermata on leukocytes population from tumor-bearing mice. (A) FATP4 mRNA expression in spleen MDSCs of LLC tumor-bearing mice treated with vehicle or lipofermata for 2 weeks evaluated via RT-qPCR analysis. (B) Percentage of MDSCs in the spleens and tumor of B16F10 tumor-bearing mice treated with or without lipofermata for 2 weeks. (C-D) Proportion of CD3^+^ T cells (C), CD4^+^ T and CD8^+^ T cells subsets among CD3^+^ T cells (D) in the spleens of B16F10 tumor-bearing mice treated with or without lipofermata for 2 weeks. (E) Lipid accumulation in MDSCs from spleens and tumor tissues of B16F10 tumor-bearing mice treated with vehicle or lipofermata for 2 weeks as determined by flow cytometry analysis. Tumor MDSCs were CD45^+^CD11b^+^Gr-1^+^ cells; For spleens: MDSCs were CD11b^+^Gr-1^+^ cells; CD3^+^T cells were CD3^+^ cells; CD4^+^ T cells were CD3^+^CD4^+^CD8^−^ cells; CD8^+^ T cells were CD3^+^CD4^−^CD8^+^ cells. Data are shown as representative of mean ± SEM in triplicate samples from two independent experiments. Unpaired student’s t-test: *p < 0.05, ns, no significant difference.

**Figure S5:**
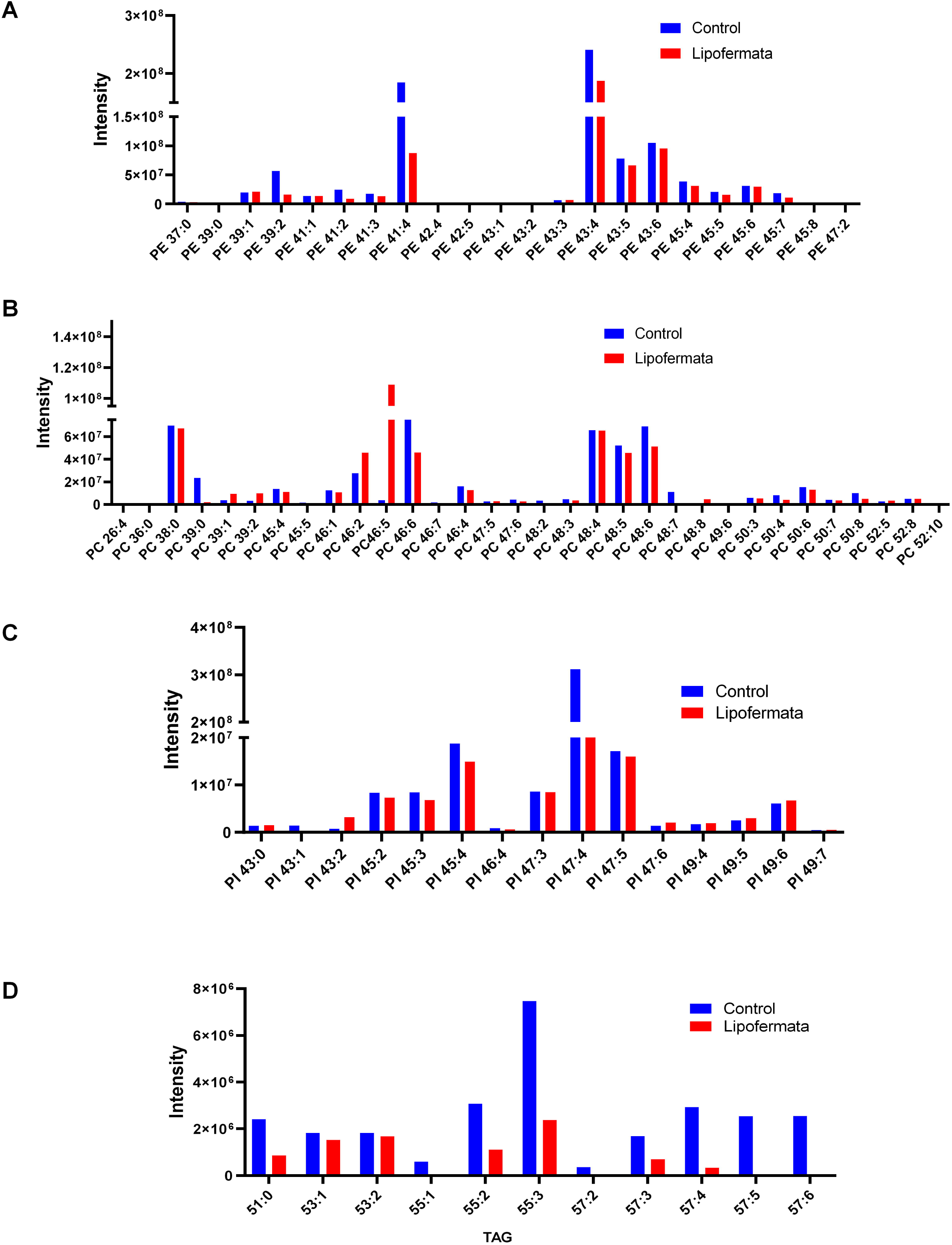
Profile of lipid molecular species in MDSCs from tumor-bearing treated with FATP2 inhibitor. (A-D) LC-MS lipid analysis for flow cytometry sorted spleen CD11b^+^ Gr1^+^ MDSCs of B16F10 tumor-bearing mice treated with or without lipofermata. Lipids extracted from MDSCs were subjected to LC-ESI-MS. Lipid molecular species were detected PE (A), PC (B), PI (C) and TAG (D). Data represents means ± SEM of relative intensity.

**Figure S6:**
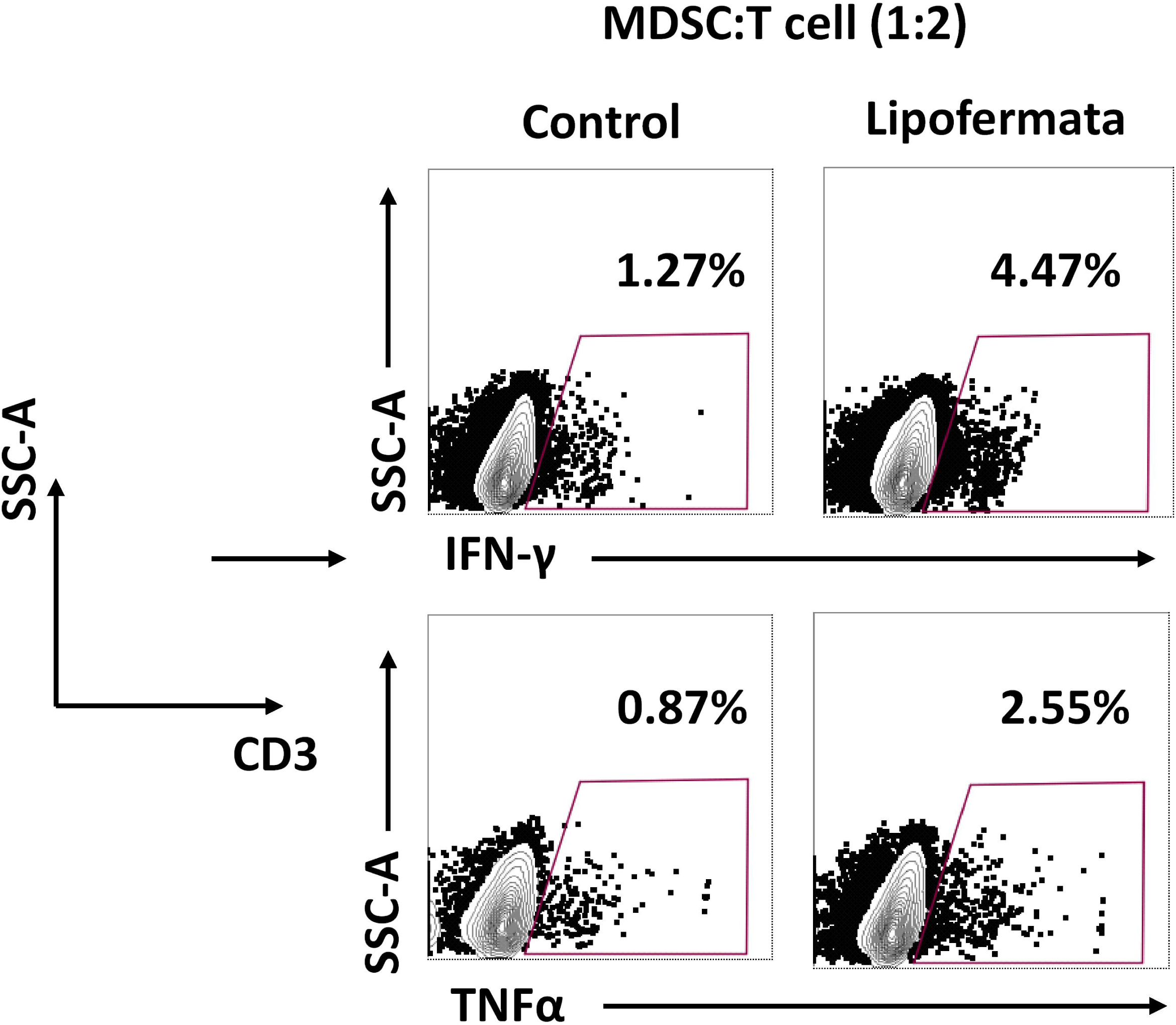
FATP2 blockade of MDSCs promotes the activation of T cells from tumor-bearing mice. Representative flow cytometry result for IFN-γ^+^ and TNFα^+^ T cells in the co-culture of spleen CD11b^+^Gr-1^+^ MDSCs and CD3^+^ T cells at ratio 1:2, respectively. MDSCs isolated from B16F10-bearing mice administered control (vehicle) or lipofermata for 2 weeks. CD3^+^ T cells were activated with the anti-CD3/CD28 purified antibody for 36 hours before testing.

**Figure S7:**
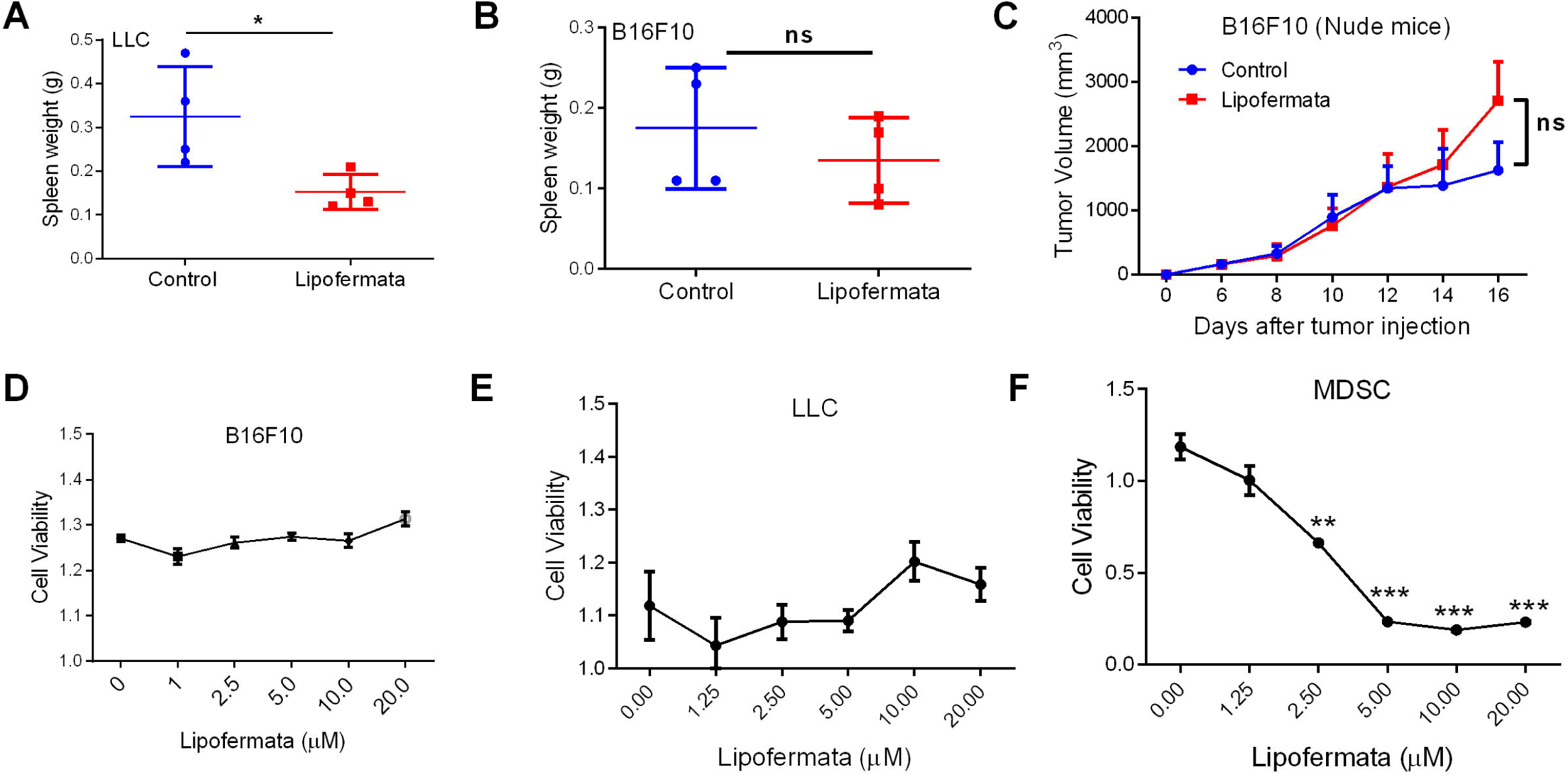
Pharmaceutical blockade of FATP2 mediated anti-tumor effect via immune mechanism. (A-B) Spleen weight from LLC (A) and B16F10 (B) tumor-bearing mice (n=4 mice/group) treated with vehicle or lipofermata for 2 weeks on days 26 and 20 post-tumor injections, respectively. (C) Tumor growth kinetics for subcutaneously injected B16F10-bearing Balb/C nude mice (n=7) treated with vehicle and lipofermata starting from day 6 when tumor size became palpable. (D-F) Cell viability assay in B16F10 cells (D), LLC cells (E) and CD11b^+^Gr1^+^ MDSCs derived from bone-marrow of B16F10 tumor-bearing mice (F) treated with the indicated concentrations of lipofermata for 72 hours. Data represents means ± SEM of two independent experiments. Unpaired student’s t-test: *p < 0.05, **p < 0.01 ***p < 0.001, ns, no significant difference.

**Figure S8:**
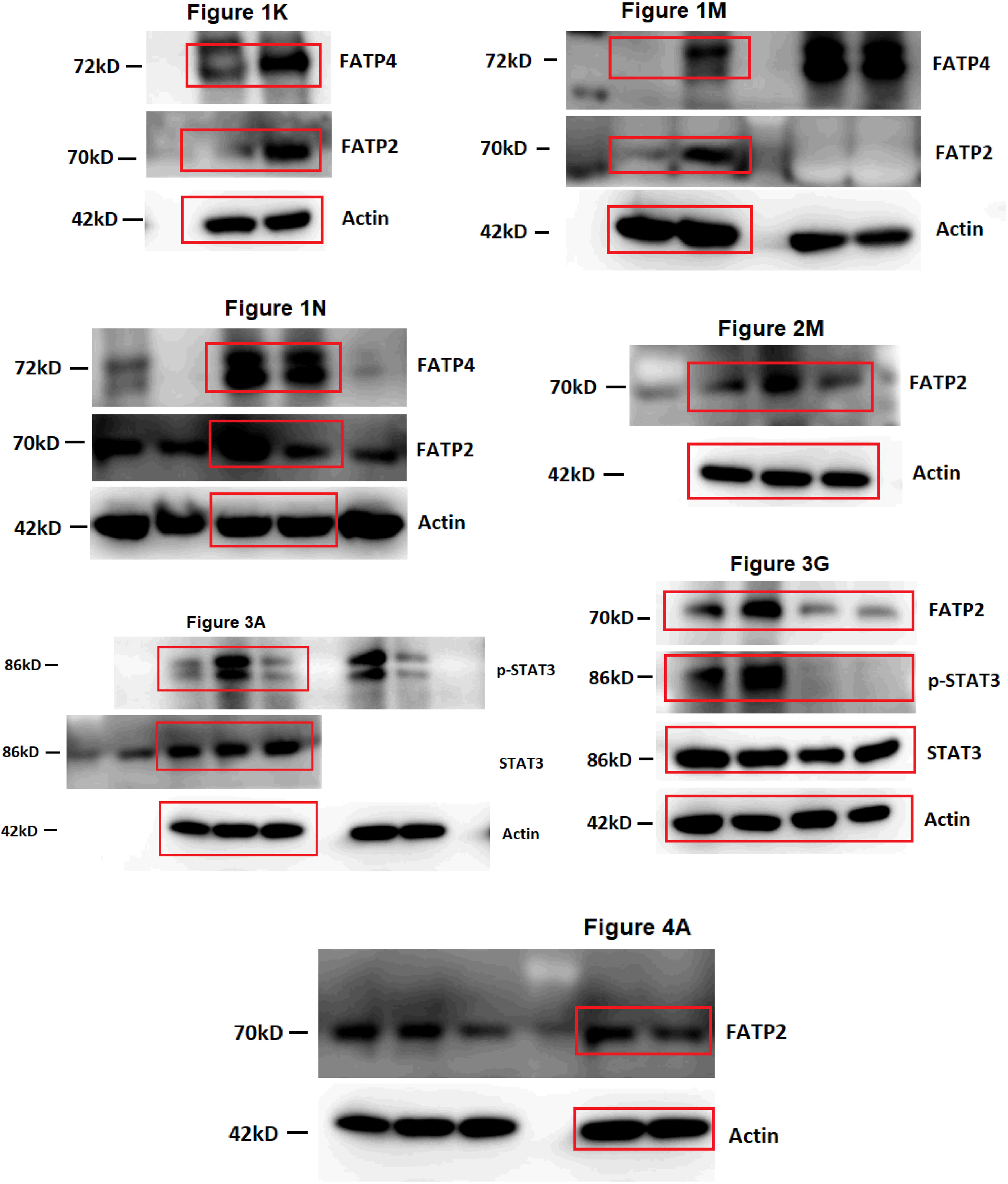
Uncropped images for western blot.

## Supplementary Tables

**Table S1:**
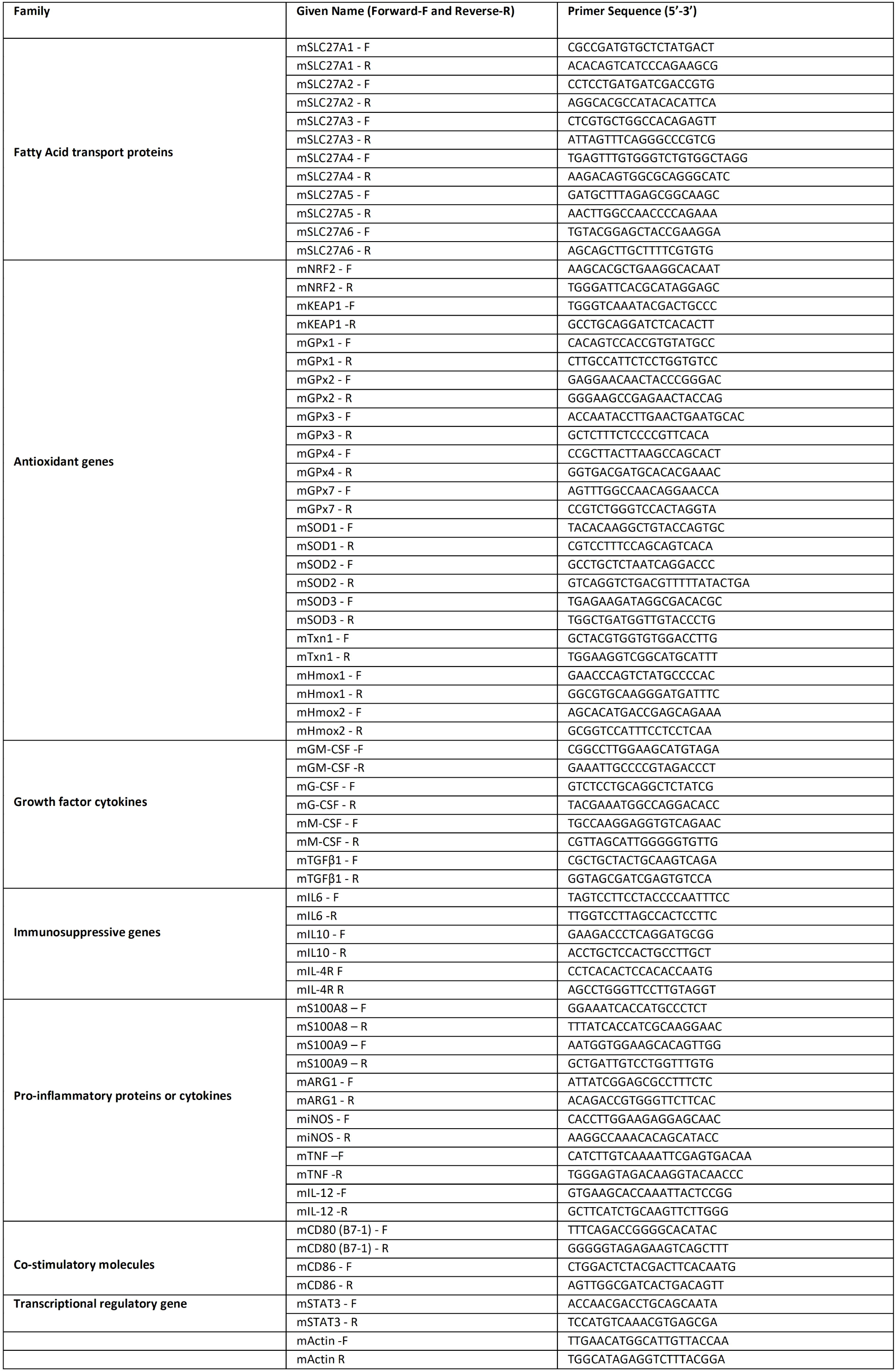
List of qPCR primer sequences

**Table S2:**
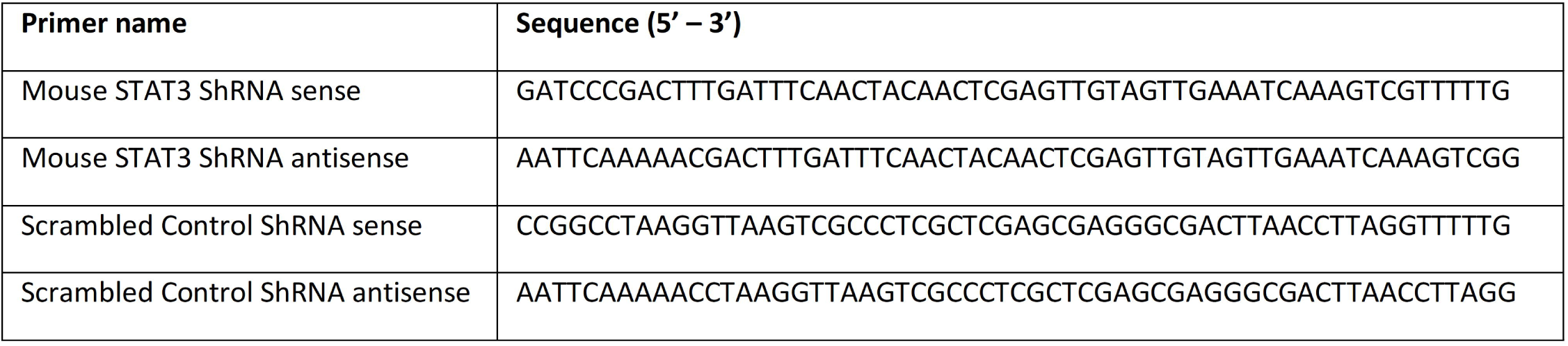
Sequences used for plasmid construction

